# A computational investigation of pathways for *ex vivo* differentiation and dysfunction of healthy and patient T cells

**DOI:** 10.64898/2026.10.26.761778

**Authors:** M. S. Lacy, Y. Binenbaum, P. R. Buenzli, A. L. Jenner

## Abstract

Adoptive T cell therapy is a promising field in cancer immunotherapy that leverages the natural ability of T cells to target and kill cancer cells. A crucial step in this therapy involves the *ex vivo* expansion of memory-like and effector-like T cells, but limiting their differentiation into exhaustion-associated phenotypes remains a challenge. Mathematical model-informed strategies to optimise T cell expansion may be able to exploit our understanding of T cell differentiation, but there is a large amount of uncertainty and disagreement in the biological literature regarding the ways in which T cells differentiate and become exhausted. We therefore conduct a large-scale mathematical investigation to help further our understanding of T cell differentiation during short-term *ex vivo* expansion. Using observations from biological literature, we first construct thousands of potential pathways connecting distinct T cell phenotypes and then formulate corresponding ordinary differential equations to test hypotheses regarding T cell differentiation and dysfunction. We fit each model to experimental data describing the *ex vivo* expansion of T cells originating from healthy individuals and cancer patients and accept those that most closely describe the data. From our findings, we hypothesise that cancer may alter T cell differentiation kinetics rather than the structure of the differentiation pathway, which has significant potential to inform therapeutic strategies. Our analysis proposes that T cells may not differentiate unidirectionally *ex vivo*; most accepted models suggest that the cells dynamically switch between functional phenotypes. We postulate that effector-like cells may be prone to upregulating inhibitory receptors that drive exhaustion, and find that modelling effector-like proliferation is unnecessary to capture observed population dynamics. Our hypothesis-testing framework generates recommendations for promising future research avenues and may pave the way to improve the production of patient-tailored T cell populations for adoptive cell therapy.

**Author summary:** T cells are immune cells that are capable of detecting and killing cancer cells, however, they are often unable to control tumour growth. Most adoptive cell therapies consider treating cancers by growing T cell populations in culture dishes outside the body and re-infusing these cells into cancer patients. T cells may be found in many unique states with varying utility in adoptive cell therapy, but it is not well understood how they transition between these states during cell culture. We propose a computational method that has the capability to address this issue. We first construct thousands of literature-informed mathematical model variations that describe the growth of relevant T cell subpopulations and compare their solutions to biological experiments. Selecting the models that best match the data, we propose hypotheses regarding characteristic pathways that connect these states, which may help to improve the production of desired T cell subtypes. Our findings also indicate that T cells may follow the same pathway to transition between states regardless of whether they originate from healthy individuals or from cancer patients. Our method may lead to computer-based optimisation strategies that could determine patient-specific culture conditions that improve adoptive cell therapy for treating cancer.

## Introduction

T cell differentiation is a crucial developmental process involved in adaptive immune responses. During an immune response, T cells undergo a series of fate decisions that commit them to differentiated and dysfunctional phenotypes [1–3]. However, the structure of the developmental pathway that connects distinct T cell phenotypes or fates—specifically memory-like, effector-like and exhausted T cells—remains unclear and is a subject of controversy [2, 4–8]. The biological literature presents conflicting ideas regarding T cell differentiation, and it is unclear whether all T cells follow intrinsic principles of differentiation or whether disease alters their developmental trajectory [9–11], making it difficult to establish a unified framework. Understanding the structure of the pathway for T cell differentiation and exhaustion will help the development of new strategies to enhance the efficacy of cancer therapies [2]. However, it is currently unclear how to prioritise biological research to reduce uncertainty. We therefore introduce a computational framework to test biological hypotheses regarding the structure of the T cell differentiation pathway during *ex vivo* expansion and make suggestions for future research avenues.

Adoptive cell therapy is a promising field in immunotherapy which often requires the *ex vivo* expansion of functional antigen-specific T cells from an initial population of tumour-infiltrating lymphocytes or T cells genetically engineered with chimeric antigen receptors (CARs) to react to tumour antigens [12–16]. Although T cells of different phenotypes play unique and important roles in immunity, not all are equally desired for applications in adoptive cell therapy [17–21]. Memory-like T cells are preferred for their ability to persist and replenish the pool of effector-like T cells [2, 22–26], but cells associated with exhaustion, which we refer to as dysfunction in this work, are known to compromise therapeutic efforts [2, 27–29]. Zhang *et al*. [30] conduct experiments in which CAR T cells are expanded using micro-rods that are designed to secrete interleukin-2 that supports T cell proliferation and present non-specific stimulatory molecules (anti-CD3 and anti-CD28) that activate cells polyclonally [31–33]. They find that varying the density of stimulus on the micro-rods has the potential to guide T cell populations towards less differentiated phenotypes with reduced expression of inhibitory receptors associated with dysfunction. This translates to superior T cell-mediated control of tumours *in vivo*, which presents promising opportunities to optimise the administration of adoptive cell therapy. However, it is still unclear how the degree of stimulation and culture time affects the yield of different T cell phenotypes and how this depends on the initial T cell population [30, 31, 34–36].

Mathematical modelling can be used for quantitative hypothesis testing of T cell differentiation and expansion [37–51]. For example, Kohler [40] uses an ordinary differential equation model to investigate the validity of a linear differentiation pathway, and Cho *et al*. [44] compare the predictions of stochastic Markov processes for two potential differentiation pathways. Computational methods have been used to simulate and understand molecular signalling involved in T cell exhaustion [52] and determine the structure of other ecological and biological networks using model selection methods [53–55]. Buchholz *et al*. [46] simulate 304 candidate pathways for *in vivo* T cell differentiation and identify two most likely ones in comparison to data. There are currently no exhaustive investigations into the structure of the pathway that connects differentiated and dysfunction-associated T cell phenotypes during *ex vivo* expansion.

In this work, we develop a quantitative hypothesis-testing framework to provide insights into T cell differentiation and dysfunction in the context of short-term *ex vivo* expansion of patient and healthy donor CAR T cells with a focus on the effects of T cell receptor stimulation. We first review the literature on relevant aspects of T cell biology and then present an extensive collection of computational models, each simulating a unique differentiation pathway. We use ordinary differential equations to model the evolution of naïve, memory-like, effector-like and dysfunction-associated cell subpopulations, and compare their predictions to T cell expansion experiments by Zhang *et al*. [30]. By investigating the most accurate mathematical models, we provide insights into T cell differentiation dynamics that are otherwise difficult to experimentally measure. We conclude by proposing promising biological research avenues that are likely to reduce uncertainty in our findings.

### T cell biology background

Many distinct phenotypes of T cells contribute to immune responses and are found during *ex vivo* expansion, namely naïve, memory stem, central memory, effector memory, and terminal effector cells [22, 25, 30]. In this work, we consider grouping T cells into naïve, memory-like and effector-like phenotypic compartments. The memory-like cell compartment contains memory stem and central memory cells. Although effector memory cells are conventionally classified as memory cells, they exhibit a more differentiated, effector-like phenotype [3, 26]. Therefore, for the purposes of our modelling framework, we classify them as effector-like cells alongside terminal effector cells. Effector-like T cells are capable of recognising and killing cells, and memory-like cells proliferate rapidly and support long-lasting immune responses by replenishing the pool of effector-like cells [3, 25]. Due to these qualities, especially in the context of cancer immunotherapy, conserving the population of functional memory T cells is critical to therapeutic success [2, 22–26].

To protect against autoreactivity and limit the immune response during chronic infections or cancer, some T cells are naturally driven towards exhausted, or dysfunctional, states upon consistent stimulation of inhibitory receptors that are upregulated after activation. Dysfunctional cells are often characterised by their minimal effector and proliferative capacities [8, 27, 56–58]. Although some tumour-infiltrating lymphocytes can recognise cancer antigens, they are often terminally differentiated or dysfunctional which complicates the expansion of less differentiated cells for therapeutic applications [2, 3, 13, 59, 60]. Late dysfunction is thought to be irreversible, but recent investigations have revealed that cells in the early stages of dysfunction may have the capabilities for rapid self-renewal and reversion to effector-like states, playing a beneficial role for immunotherapies [8, 29, 56, 61].

Table 1 summarises the markers that we use to define T cell compartments in our computational framework, which are designed to provide a clear way to distinguish T cell subpopulations and relate our models to experimental data. We use the cell markers CD45RA and CCR7 to describe the differentiation state of T cells. CD95 is a marker that indicates stemness, so naïve cells are often identified from their lack of CD95 expression [62]. In the case where CD95 expression data is not available, we also assume that they may be identified by a lack of expression of CD25, which is a marker that indicates activation [63]. We model a subpopulation of expanded cells upregulating the inhibitory receptors PD-1 and TIM-3, which may be dysfunctional or prone to becoming dysfunctional following adoptive transfer [56]. We refer to cells co-expressing these inhibitory receptors as dysfunction-associated.

**Table 1.** Marker expressions to model T cell subtypes. Expressions that we use to identify T cell phenotypic compartments, where + indicates that the modelled subtype expresses the marker, and – indicates that it does not. In the PD-1/TIM-3 column, the first marker indicates PD-1 expression and the second indicates TIM-3 expression. Blank cells indicate that the marker was not considered (may be + or –).

| T cell subtype | CD45RA | CCR7 | CD95 | CD25 | PD-1/TIM-3 | Reference |
| --- | --- | --- | --- | --- | --- | --- |
| Naïve | + | + | – | – | –/–, +/–, or –/+ | [62, 63] |
| Memory stem (memory-like) | + | + | + | + | –/–, +/–, or –/+ | [62] |
| Central memory (memory-like) | – | + |  |  | –/–, +/–, or –/+ | [62] |
| Effector memory (effector-like) | – | – |  |  | –/–, +/–, or –/+ | [62] |
| Terminal effector (effector-like) | + | – |  |  | –/–, +/–, or –/+ | [62] |
| Dysfunction-associated |  |  |  |  | +/+ | [56] |

The biological literature proposes many potential pathways for differentiation between memory-like, effector-like and dysfunctional T cells, some involving fate decisions that differ between acute and chronic antigen stimulation [3, 7, 8, 11, 27, 28, 56, 64–66]. Some studies suggest that T cells differentiate in a linear fashion [23, 25, 67–69], while others propose branching pathways [8, 68–71]. Bidirectional exchanges between memory-like and effector-like T cells are also suggested, often in the context of *in vivo* immune responses [2, 4, 5, 72]. Effector-like cells are often thought to be precursors to dysfunction, but it is also suggested that naïve and memory-like cells may become dysfunctional [8, 56–58, 64, 65, 73]. It is currently unclear whether individual naïve cells commit to a predetermined fate or whether T cells have the capacity for plastic, or adaptive, differentiation [6, 10, 74, 75]. For the purpose of mathematical modelling, we consider that differentiation pathways may be deconstructed into features of differentiation, dysfunction, proliferation, and death. Table 2 presents a comprehensive list of commonly referenced features of the T cell differentiation pathway, and our interpretation of the support for each feature in the biological literature in the context of short-term *ex vivo* expansion (detailed further in S1 Text).

**Table 2.**
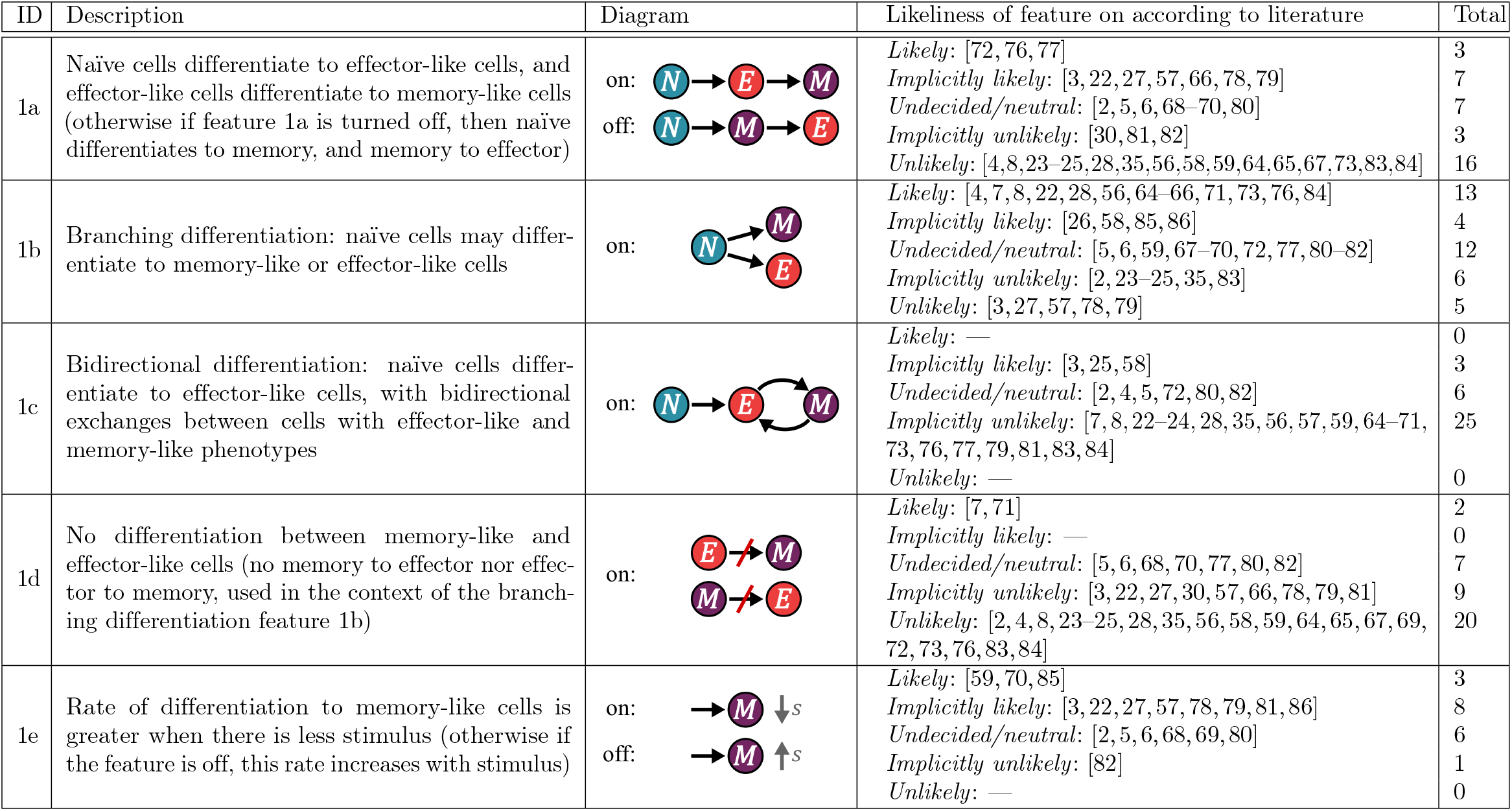
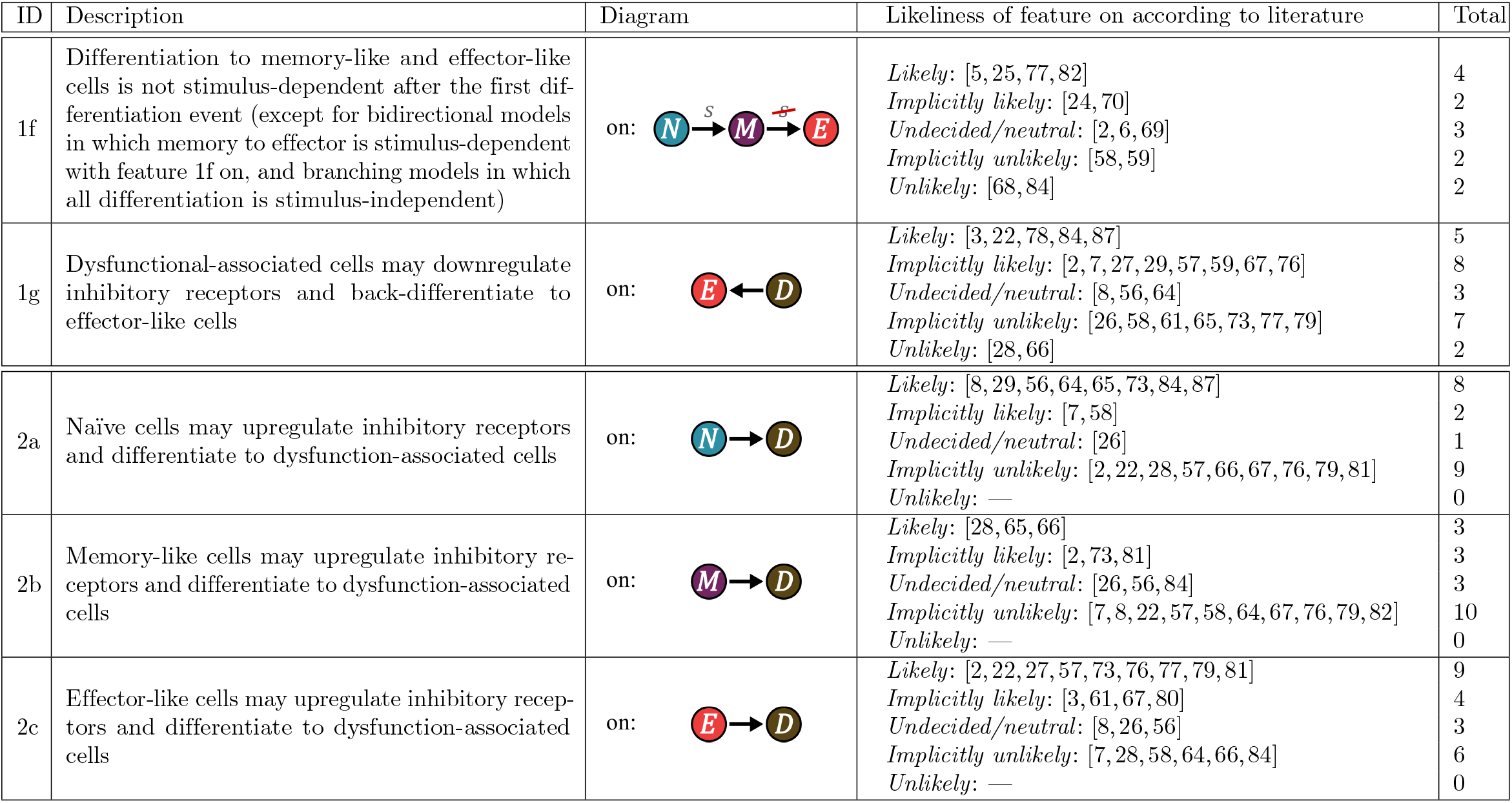
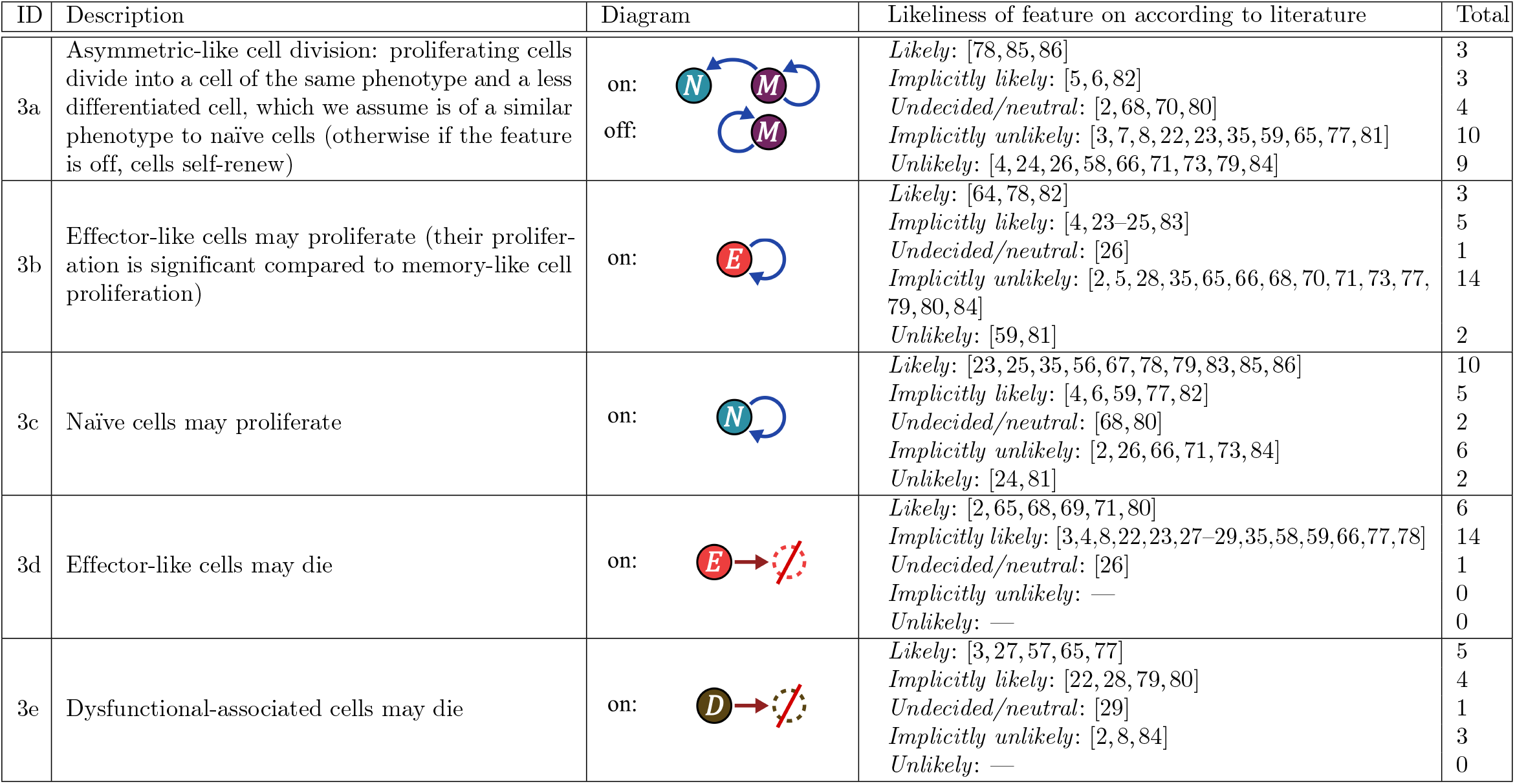
Literature review of T cell differentiation features. This table reviews features of T cell differentiation pathways proposed in the biological literature. Features 1a–1g relate to differentiation, 2a–2c relate to the upregulation of inhibitory receptors associated with dysfunction, and 3a–3e relate to proliferation and death. The diagrams show example pathway structures that may be generated when the feature is turned on or off. In the diagrams, *N* refers to naïve or less differentiated cells (blue), *M* refers to memory-like cells (purple), *E* refers to effector-like cells (red), *D* refers to dysfunction-associated cells (brown), and *s* refers to the presence of stimulatory molecules. Black arrows indicate differentiation (or regulation of inhibitory receptors in the context of dysfunction-associated cells), blue arrows indicate cell proliferation, and red arrows indicate cell death. We present the likeliness of each feature to be on during *ex vivo* T cell expansion according to our interpretation of the literature, alongside the total number of articles cited with the same likeliness.

| ID | Description | Diagram | Likelihood of feature on according to literature | Total |
| --- | --- | --- | --- | --- |
| 1a | Naïve cells differentiate to effector-like cells, and effector-like cells differentiate to memory-like cells (otherwise if feature 1a is turned off, then naïve differentiates to memory, and memory to effector) | on:<br>off: | *Likely*: [72, 76, 77]<br>*Implicitly likely*: [3, 22, 27, 57, 66, 78, 79]<br>*Undecided/neutral*: [2, 5, 6, 68–70, 80]<br>*Implicitly unlikely*: [30, 81, 82]<br>*Unlikely*: [4, 8, 23–25, 28, 35, 56, 58, 59, 64, 65, 67, 73, 83, 84] | 3<br>7<br>7<br>3<br>16 |
| 1b | Branching differentiation: naïve cells may differentiate to memory-like or effector-like cells | on: | *Likely*: [4, 7, 8, 22, 28, 56, 64–66, 71, 73, 76, 84]<br>*Implicitly likely*: [26, 58, 85, 86]<br>*Undecided/neutral*: [5, 6, 59, 67–70, 72, 77, 80–82]<br>*Implicitly unlikely*: [2, 23–25, 35, 83]<br>*Unlikely*: [3, 27, 57, 78, 79] | 13<br>4<br>12<br>6<br>5 |
| 1c | Bidirectional differentiation: naïve cells differentiate to effector-like cells, with bidirectional exchanges between cells with effector-like and memory-like phenotypes | on: | *Likely*: —<br>*Implicitly likely*: [3, 25, 58]<br>*Undecided/neutral*: [2, 4, 5, 72, 80, 82]<br>*Implicitly unlikely*: [7, 8, 22–24, 28, 35, 56, 57, 59, 64–71, 73, 76, 77, 79, 81, 83, 84]<br>*Unlikely*: — | 0<br>3<br>6<br>25<br>0 |
| 1d | No differentiation between memory-like and effector-like cells (no memory to effector nor effector to memory, used in the context of the branching differentiation feature 1b) | on: | *Likely*: [7, 71]<br>*Implicitly likely*: —<br>*Undecided/neutral*: [5, 6, 68, 70, 77, 80, 82]<br>*Implicitly unlikely*: [3, 22, 27, 30, 57, 66, 78, 79, 81]<br>*Unlikely*: [2, 4, 8, 23–25, 28, 35, 56, 58, 59, 64, 65, 67, 69, 72, 73, 76, 83, 84] | 2<br>0<br>7<br>9<br>20 |
| 1e | Rate of differentiation to memory-like cells is greater when there is less stimulus (otherwise if the feature is off, this rate increases with stimulus) | on:<br>off: | *Likely*: [59, 70, 85]<br>*Implicitly likely*: [3, 22, 27, 57, 78, 79, 81, 86]<br>*Undecided/neutral*: [2, 5, 6, 68, 69, 80]<br>*Implicitly unlikely*: [82]<br>*Unlikely*: — | 3<br>8<br>6<br>1<br>0 |
| 1f | Differentiation to memory-like and effector-like cells is not stimulus-dependent after the first differentiation event (except for bidirectional models in which memory to effector is stimulus-dependent with feature 1f on, and branching models in which all differentiation is stimulus-independent) | on: | *Likely*: [5, 25, 77, 82]<br>*Implicitly likely*: [24, 70]<br>*Undecided/neutral*: [2, 6, 69]<br>*Implicitly unlikely*: [58, 59]<br>*Unlikely*: [68, 84] | 4<br>2<br>3<br>2<br>2 |
| 1g | Dysfunctional-associated cells may downregulate inhibitory receptors and back-differentiate to effector-like cells | on: | *Likely*: [3, 22, 78, 84, 87]<br>*Implicitly likely*: [2, 7, 27, 29, 57, 59, 67, 76]<br>*Undecided/neutral*: [8, 56, 64]<br>*Implicitly unlikely*: [26, 58, 61, 65, 73, 77, 79]<br>*Unlikely*: [28, 66] | 5<br>8<br>3<br>7<br>2 |
| 2a | Naïve cells may upregulate inhibitory receptors and differentiate to dysfunction-associated cells | on: | *Likely*: [8, 29, 56, 64, 65, 73, 84, 87]<br>*Implicitly likely*: [7, 58]<br>*Undecided/neutral*: [26]<br>*Implicitly unlikely*: [2, 22, 28, 57, 66, 67, 76, 79, 81]<br>*Unlikely*: — | 8<br>2<br>1<br>9<br>0 |
| 2b | Memory-like cells may upregulate inhibitory receptors and differentiate to dysfunction-associated cells | on: | *Likely*: [28, 65, 66]<br>*Implicitly likely*: [2, 73, 81]<br>*Undecided/neutral*: [26, 56, 84]<br>*Implicitly unlikely*: [7, 8, 22, 57, 58, 64, 67, 76, 79, 82]<br>*Unlikely*: — | 3<br>3<br>3<br>10<br>0 |
| 2c | Effector-like cells may upregulate inhibitory receptors and differentiate to dysfunction-associated cells | on: | *Likely*: [2, 22, 27, 57, 73, 76, 77, 79, 81]<br>*Implicitly likely*: [3, 61, 67, 80]<br>*Undecided/neutral*: [8, 26, 56]<br>*Implicitly unlikely*: [7, 28, 58, 64, 66, 84]<br>*Unlikely*: — | 9<br>4<br>3<br>6<br>0 |
| 3a | Asymmetric-like cell division: proliferating cells divide into a cell of the same phenotype and a less differentiated cell, which we assume is of a similar phenotype to naïve cells (otherwise if the feature is off, cells self-renew) | on:<br>off: | *Likely*: [78, 85, 86]<br>*Implicitly likely*: [5, 6, 82]<br>*Undecided/neutral*: [2, 68, 70, 80]<br>*Implicitly unlikely*: [3, 7, 8, 22, 23, 35, 59, 65, 77, 81]<br>*Unlikely*: [4, 24, 26, 58, 66, 71, 73, 79, 84] | 3<br>3<br>4<br>10<br>9 |
| 3b | Effector-like cells may proliferate (their proliferation is significant compared to memory-like cell proliferation) | on: | *Likely*: [64, 78, 82]<br>*Implicitly likely*: [4, 23–25, 83]<br>*Undecided/neutral*: [26]<br>*Implicitly unlikely*: [2, 5, 28, 35, 65, 66, 68, 70, 71, 73, 77, 79, 80, 84]<br>*Unlikely*: [59, 81] | 3<br>5<br>1<br>14<br>2 |
| 3c | Naïve cells may proliferate | on: | *Likely*: [23, 25, 35, 56, 67, 78, 79, 83, 85, 86]<br>*Implicitly likely*: [4, 6, 59, 77, 82]<br>*Undecided/neutral*: [68, 80]<br>*Implicitly unlikely*: [2, 26, 66, 71, 73, 84]<br>*Unlikely*: [24, 81] | 10<br>5<br>2<br>6<br>2 |
| 3d | Effector-like cells may die | on: | *Likely*: [2, 65, 68, 69, 71, 80]<br>*Implicitly likely*: [3, 4, 8, 22, 23, 27–29, 35, 58, 59, 66, 77, 78]<br>*Undecided/neutral*: [26]<br>*Implicitly unlikely*: —<br>*Unlikely*: — | 6<br>14<br>1<br>0<br>0 |
| 3e | Dysfunctional-associated cells may die | on: | *Likely*: [3, 27, 57, 65, 77]<br>*Implicitly likely*: [22, 28, 79, 80]<br>*Undecided/neutral*: [29]<br>*Implicitly unlikely*: [2, 8, 84]<br>*Unlikely*: — | 5<br>4<br>1<br>3<br>0 |

## Methods

### Experimental data

Zhang *et al*. [30] consider the short-term *ex vivo* expansion of CAR T cells derived from healthy individuals and patients with acute lymphoblastic leukemia and chronic lymphocytic lymphoma. They reveal the proportions of cells that express markers associated with differentiation and dysfunction after eight days of *ex vivo* expansion with interleukin-2 supplementation and under varying densities of anti-CD3 and anti-CD28 (at an anti-CD3-to-anti-CD28 ratio of 1:1) attached to lipids coating the micro-rods. Their data indicates that the level of polyclonal stimulation may impact T cell differentiation, which has also been demonstrated in another study by Shi *et al*. [88]. These experiments do not describe the effects of varying the concentrations of interleukin-2 or other cytokines on the expanded T cell subpopulations. Findings from other studies indicate that the use of interleukin-2 may have driven T cell differentiation towards cells with effector-like and dysfunction-associated phenotypes [66, 68, 69, 89]. Using the definitions in Table 1, we compute the numbers of naïve, memory-like, effector-like and dysfunction-associated cells at days 0 and 8 for each experiment. Our calculations and assumptions are detailed in S2 Text.

### Mathematical model

We consider the density of stimulus attached to the surfaces of micro-rods as a key variable in our modelling framework as differentiation and dysfunction seem to be driven by stimulation of the T cell receptor complex [30]. To reduce model complexity, we group T cell states into four model compartments (seen in Fig 1) with phenotypes defined by the markers in Table 1:

**Fig 1.**
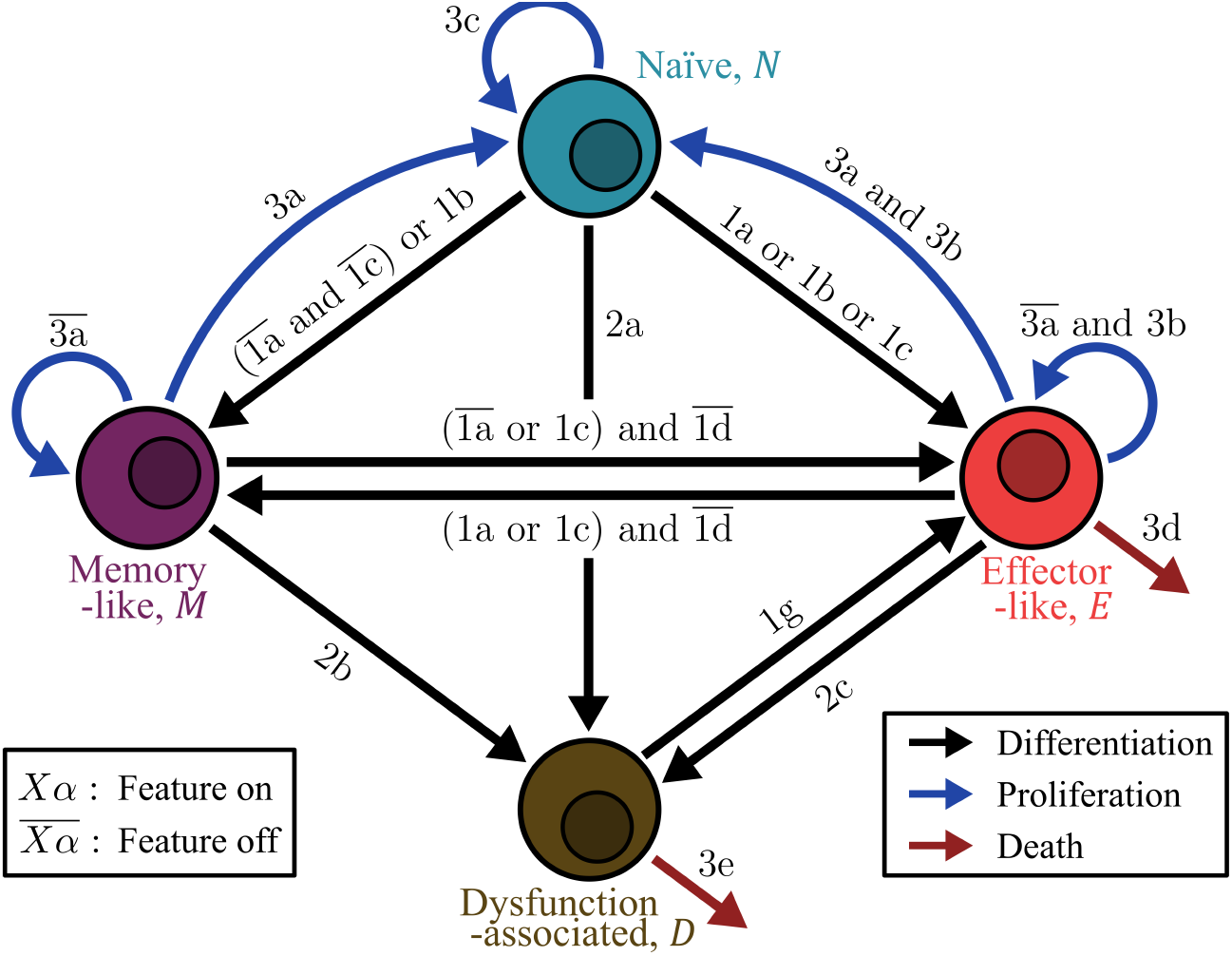
Diagram of all pathways simulated by mathematical models. Diagram depicting the structures of modelled differentiation pathways between naïve *N*, memory-like *M*, effector-like *E*, and dysfunction-associated *D* cells. These pathways are conditional on the selections made for literature-informed features in Table 2 of the form *Xα* (*X* ∈ {1, 2, 3} and *α* ∈ {a, b, c, …}). Features 1e and 1f are not represented in this diagram as they only affect the rates of differentiation, not the pathway structure.

1. Naïve cells *N* —less differentiated T cells that have not yet been activated. We assume that these cells do not die within eight days regardless of feature selections, as this is indicated by most articles cited in Table 2.
2. Memory-like cells *M* —activated memory stem and central memory T cells. We assume that these cells proliferate and do not die within eight days, as indicated by most articles in Table 2.
3. Effector-like cells *E*—activated effector memory and terminal effector T cells.
4. Dysfunction-associated cells *D*—T cells co-expressing inhibitory receptors PD-1 and TIM-3. We assume that some of these cells exhibit characteristics similar to late dysfunctional cells and therefore omit dysfunction-associated cell proliferation from our model.

For model simplicity and due to a lack of data regarding dysfunction markers on distinct T cell phenotypes, we assume that cells expressing markers for naïve, memory-like and effector-like phenotypes are dysfunction-associated cells if they also co-express inhibitory receptors PD-1 and TIM-3 (see Table 1). We treat T cells expressing CD4 and CD8 equivalently in this framework.

To capture uncertainty in our current understanding of T cell differentiation, we construct various potential T cell differentiation pathways exhibiting the literature-informed features in Table 2. Each feature may be selectively turned on or off which adds or removes fate decisions from the pathway, except for features 1e and 1f which only alter rates of differentiation. Fig 1 illustrates all pathway structures that can be created using combinations of feature selections. For example, turning feature 1a on may describe linear differentiation from naïve to effector-like and then to memory-like states, but turning it off (denoted by a bar above 1a, i.e. 1a) switches differentiation to proceed from naïve to memory-like to effector-like states. A branching model is created by turning feature 1b on, in which naïve cells may transition to either the memory-like or effector-like compartments. Turning both features 1b and 1a on constructs a differentiation pathway which branches from naïve to memory-like or effector-like states, and includes a transition from effector-like to memory-like cells, which is proposed by some articles in the literature [66, 76]. If feature 1d was turned on in a model with feature 1b on, it would describe a branching model without differentiation between memory-like and effector-like states, which is suggested in other works [7, 71].

Each unique combination of feature selections (on or off) represents a pathway with a unique structure, and most pathways proposed by articles cited in Table 2 may be estimated by one of our models. Accounting for excluded feature combinations described in S3 Text, this method allows for 7,840 unique T cell differentiation pathways between naïve, memory-like, effector-like and dysfunction-associated phenotypic compartments. To generate a mathematical model that simulates one of these pathways, we assume that the time evolution of T cell subpopulations may be described on a macroscopic scale with in-going sources and out-going sinks for each cell type. We use first-order kinetic rates, which are modulated by the molar mass ratio of stimulatory molecules (anti-CD3 and anti-CD28) to lipids coating the micro-rods *s*. These include rates of differentiation to a memory-like state *r*_m_(*s*), differentiation to an effector-like state *r*_e_(*s*), and upregulation of dysfunction-associated markers *r*_d_(*s*), which do not change over time since the stimulus density *s* is time-invariant in our model. We also introduce rates of cell proliferation *µ* and cell death *λ*, which we assume to be constant during short-term *ex vivo* expansion for simplicity.

All mathematical models are represented by a system of linear ordinary differential equations, which simulate the population dynamics of naïve cells *N* (*t, s*), memory-like cells *M* (*t, s*), effector-like cells *E*(*t, s*) and dysfunction-associated cells *D*(*t, s*) at time *t* and under stimulus density *s*:

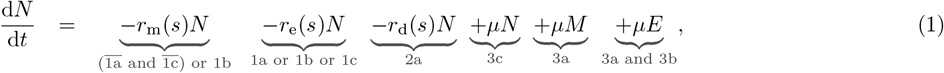

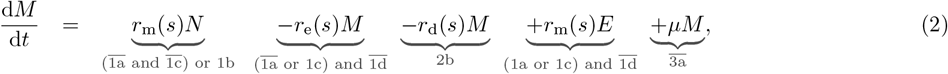

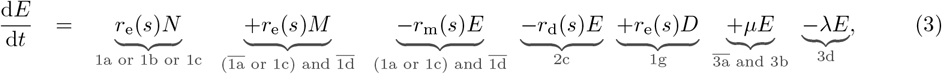

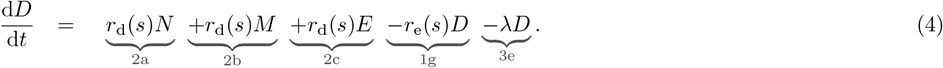

Each term in the equations is only present if the condition below the term is met (corresponding to conditions in Fig 1). For example, the first terms in the first two equations correspond to differentiation from naïve cells to memory-like cells and are only present in the model if features 1a (differentiation from naïve to effector to memory) and 1c (differentiation is bidirectional, starting with naïve to effector) are both off (represented by bars above 1a and 1c, i.e. 1a and 1c) or feature 1b (branching differentiation from naïve to effector or memory) is on. See S4 Text for details on the formulation of our mathematical model and reasoning for the conditions on pathway features.

For any individual model with a given set of parameters, the system of linear ordinary differential equations (Eqs (1)–(4)) is of the form 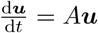, where ***u*** = [*N, M, E, D*]^T^ contains cell counts for all T cell compartments and *A* is the coefficient matrix. The solution to this system at time *t* is ***u*** = *e*^*At*^***u***_**0**_, where ***u***_**0**_ = [*N* (0), *M* (0), *E*(0), *D*(0)]^T^ contains the initial cell counts from the data and *e*^*At*^ is evaluated numerically (see S5 Text for more details). Our mathematical framework does not model spatial dynamics, such as interactions between T cells and the micro-rods presenting stimuli. We have previously shown that a non-spatial ordinary differential equation model of T cell expansion with micro-rods can emulate the population predictions of spatial models when the micro-rods are well-mixed [37], which is the case at the beginning of *ex vivo* expansion experiments with micro-rods [32].

When feature 1f is off, the stimulus dependence of the rates of differentiation and dysfunction are defined by Michaelis-Menten equations, which are commonly used to describe reaction rates as functions of the concentration of a substance [90]. The equation for the rate of differentiation to the memory-like compartment changes with feature 1e such that *r*_m_ decreases as *s* increases. With feature 1f off, these rates are, respectively,

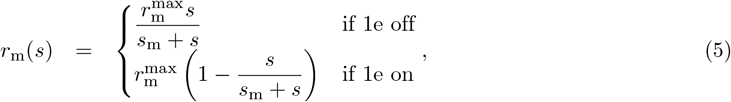

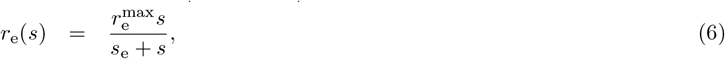

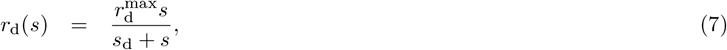

where 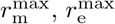 and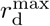 are the maximum rates of differentiation to memory-like, effector-like, and dysfunction-associated cells, respectively, and *s*_m_, *s*_e_ and *s*_d_ are the half-maximum stimulus densities encoding sensitivity to stimulus [90]. When feature 1f is on (stimulus-dependence only for the first differentiation event) and if naïve cells do not differentiate directly to memory-like cells, we assume that 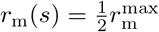 and similarly for differentiation to an effector-like state. These rates are further detailed in S4 Text.

We allow the values for these parameters alongside the proliferation rate *µ* to vary depending on the pathway structure, but for simplicity, we fix the T cell death rate. We select *λ* = 0.1 day^−1^ for both effector-like and dysfunction-associated cells, as we found this value to reasonably describe T cell population dynamics in a previous mathematical model of T cell expansion [37]. We also assume that the rates of differentiation, dysfunction and proliferation are the same for each cell type under a given stimulus density for simplicity and ease of model fitting. Under most parameter regimes, this model describes approximately exponential growth of each cell subpopulation, which seems to be a reasonable approximation of short-term *ex vivo* expansion [30, 31]. Although it has been reasoned that mathematical models of T cell differentiation should allow for these rates to also depend on the cell generation number [38], we argue that a simpler model is sufficient to capture 8-day expansion. We do not consider modelling the clonal contraction phase that occurs after expansion in a typical immune response.

### Model fitting

All model fitting and analysis is performed in Julia (v1.12.3) [91], with code available on GitHub [92].

It is expected that the structures of the T cell differentiation pathways paired with stimulus modulations influence cell population dynamics at the final time *T* = 8 days [4]. This implies that the validity of each pathway may be evaluated by comparing population predictions from the corresponding mathematical model to the stimulus-dependent data by Zhang *et al*. [30]. We consider fitting numerical solutions of Eqs (1)–(4) at Day 8 (under specific feature selections) to the average expanded healthy T cell counts at Day 8 for each stimulus density *s* by estimating the parameters 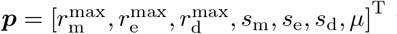 that minimise the healthy donor loss function

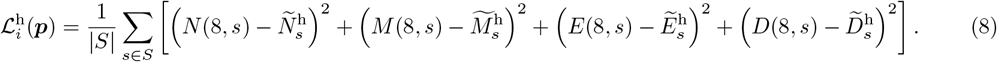

This loss describes the sum of squared errors between cell subpopulations in our *i*th model (*i* = 1, …, 7840) and the healthy data set, averaged over the stimulus densities used in experiments. Here, |*S*| = 7 is the number of stimulus densities with *S* = {0.8, 2, 4, 6, 8, 10, 12} (mass in nanograms of activating stimulus per microgram of micro-rods) and *N* ^h^ is the average number of healthy naïve cells at Day 8 from experiments with micro-rods expressing stimuli with density *s* (and similarly for the other simulated cell subtypes). We use a similar expression to represent a cancer patient loss function 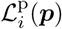 for the *i*th model based on the patient data set 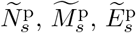, and 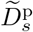.

For each of the 7,840 mathematical models describing unique differentiation pathways, we implement the Broyden–Fletcher–Goldfarb–Shanno optimisation algorithm using Optim.jl in Julia [93] to minimise the loss in Eq (8). To avoid local minima during optimisation, we implement a multi-start approach by sampling ten distinct initial parameter guesses from a bounded parameter space using Latin hypercube sampling with LatinHypercubeSampling.jl [94]. We limit the search in parameter space such that each fitted parameter is bounded below by 0 and above by biologically reasonable values. Namely, the maximal rates of differentiation and dysfunction 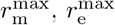 and 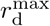 are bounded by 20 day^−1^, the stimulus densities *s*_m_, *s*_e_ and *s*_d_ are bounded by 30 ng/µg (stimulus to lipid mass ratio), and the proliferation rate *µ* is bounded by 10 day^−1^. See S5 Text for details on our methods for optimising computations during model fitting.

To compare minimal loss between models, we normalise 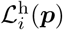 and 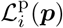 by the average minimal loss over all models:

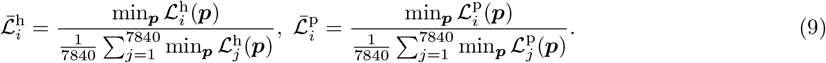

### Analysis of results

Rather than selecting a single model that most closely matches the healthy data for example, all models whose predictions sufficiently describe the healthy data are accepted to represent differentiation pathways that T cells from healthy individuals may follow during *ex vivo* expansion. However, to also take into account how well these models describe the cancer patient data, we consider the average of the normalised minimal losses for the *i*th model:

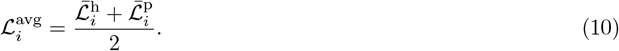

Because models with low average loss do not necessarily predict T cell expansion accurately for all data points, we accept models that have both low average loss and whose predictions lie within one standard deviation about the mean of the data (detailed further in the Results section). This enables us to account for uncertainty in experimental measurements and model calibration. We choose to not use metrics such as the Akaike information criterion which penalises models with many parameters [95] since most of our models have the same number of parameters. While it may be possible to implement methods that directly fit the structure of the differentiation pathway to data, we choose to methodically evaluate all potential pathways so that we may also uncover information regarding pathways that do not match the data precisely.

Many models that explain the data have similar features. We interpret a feature common to many well-performing models to be more likely to describe T cell differentiation in the *ex vivo* experiments. To quantify this likeliness, we use Yule’s Q coefficient of association [96]. Let *A* represent the event that a model is accepted to sufficiently capture experimental data (as defined above), and *Xα* be the event that some feature *Xα* is turned on in the model (*X* ∈ {1, 2, 3} and *α* ∈ {a, b, c, …}). Yule’s Q coefficient compares probabilities that a model is accepted if it has feature *Xα* turned on, Pr(*A* | *Xα*), to the probability that a model is accepted if it does not have the feature turned on, 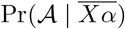. These probabilities are found with Bayes’ theorem, which uses the frequencies of each feature selection among the accepted models and among all simulated models. Yule’s Q coefficient (*Q*) is then a re-scaling of the appropriate odds ratio OR such that −1 ≤ *Q* ≤ 1;

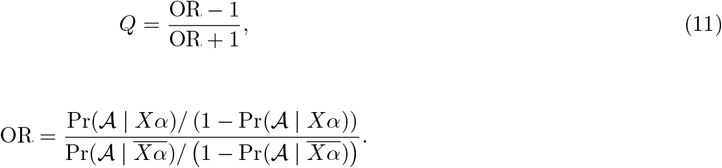

where

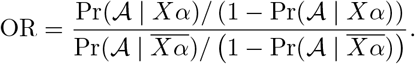

When *Q* is positive, models with the feature switched on are more likely to be accepted than models without the feature, and *Q* = 0 indicates that the feature is not associated with model acceptance. Negative values of *Q* may be used to infer the probability that a model is accepted if the feature is turned off, except for feature 1a since some models may be considered to have neither feature 1a turned on nor off (this is when features 1c or 1d are on, see S3 Text), so we evaluate Yule’s Q coefficient for both feature 1a on and off. We emphasise that, since expansion is occurring in a controlled environment with interleukin-2 supplementation and polyclonal stimulation, our results may only inform the T cell differentiation pathway for similar short-term controlled experiments.

We are also interested in comparing our predictions regarding the likeliness of differentiation features to the biological literature. We consider constructing a summary of the literature’s support for each feature as a weighted average of the total number of articles that indicate the likeliness of a feature in Table 2.

Specifically, for a given feature selection with *L*_2_ articles listed as likely, *L*_1_ as implicitly likely, *L*_0_ as neutral, *L*_−1_ as implicitly unlikely, and *L*_−2_ as unlikely, the weighted average is given as

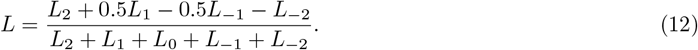

This quantity ranges from −1 to 1 and is interpreted similarly to Yule’s Q coefficient regarding the likeliness of each feature.

## Results

### Ranking healthy and patient T cell models

We first fit all 7,840 mathematical models to the data by Zhang *et al*. [30] and rank models of individual pathways against those of all other pathways based on increasing order of normalised minimum loss 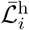 and 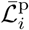 in Eq (9). Fig 2a shows the normalised minimum loss 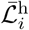 for models fitted to the healthy (left) and 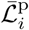 patient (right) data sets. We see that predictions from only approximately the first 500 models consistently lie within one standard deviation about the mean of cell counts for both data sets (blue). Note that some models with predictions that lie outside one standard deviation about the mean (red) have a lower loss than other models whose predictions lie within one standard deviation. We also see that the worst 2,000 models (ranking from around #5,840 to #7,840) have a substantially higher loss than the most accurate models, which suggests that these represent pathways that are unlikely to be representative of the true differentiation pathway. S1 Video and S2 Video show pathway diagrams for all models ordered by increasing healthy donor and patient losses.

**Fig 2.**
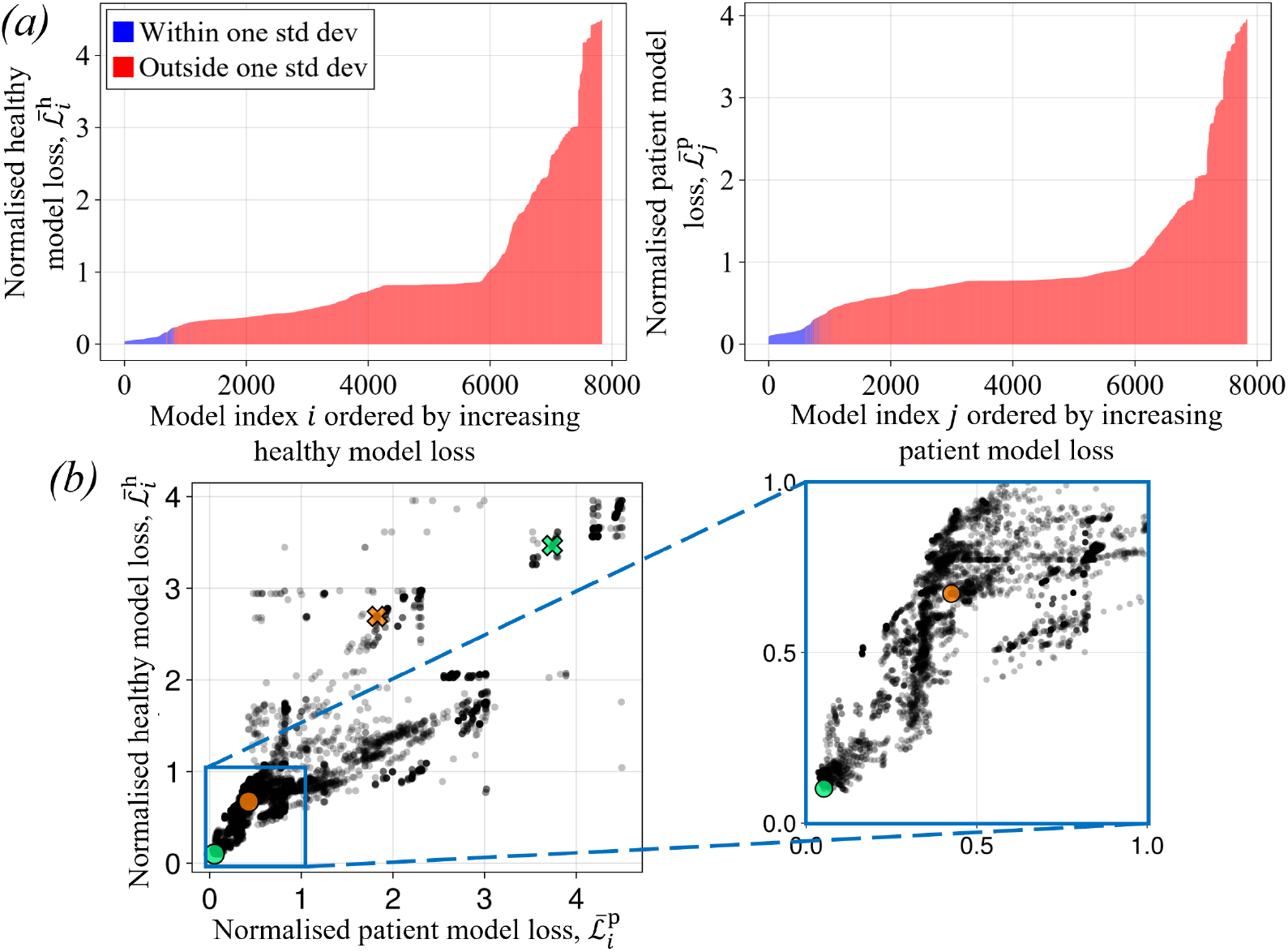
Comparison of models fitted to the healthy and patient data sets. *(a)* Normalised healthy donor (left) and patient (right) loss (Eq (9)) ordered by increasing loss, coloured by models whose Day 8 predictions lie within (blue) or outside (red) one standard deviation about the mean of the corresponding data set for all stimulus densities and all T cell compartments except naïve cells. *(b)* Normalised healthy donor and patient losses for each model (left), with a zoomed plot showing models with the lowest losses (right). Points represent individual models out of the 7,840 fitted models. The coloured crosses indicate the base models and the coloured circles indicate the altered models for the first (green, see Fig 3) and second (orange, see Fig 4) case studies.

When comparing the healthy donor and patient losses for a given model (Fig 2b), we find that most pathways tend to describe the differentiation of healthy and patient-derived T cells with similar accuracy. Under our modelling framework, healthy donor loss is correlated to patient loss with a Pearson correlation coefficient of 0.896. It is important to note here that, although one pathway structure may accurately explain both data sets, the corresponding model is fitted to each data set with different parameter values. This indicates that individual model structures of T cell differentiation are likely to yield predictions of healthy and patient T cell expansion with similar accuracy, up to recalibration in parameter values.

### Case studies on features of differentiation

Many differentiation pathways proposed in the biological literature are represented by individual models in Fig 2, some of which we explore further here. As our first case study, Wherry [57] proposes a pathway in which naïve cells transition to effector-like cells, which then transition to memory-like or dysfunctional cells. They describe a range of dysfunctional states, the earliest of which may have the potential to back-differentiate to an effector-like state, and the latest of which eventually dies. A pathway that approximates their suggestion is depicted in Fig 3a under the transitions indicated by solid arrows, which is simulated by the ordinary differential equation model in Fig 3b. To approximate their suggestion for T cell dysfunction, we initially assume that only effector-like cells can upregulate inhibitory receptors. The self-renewal of effector-like cells is not considered in this model, but it is considered by other models represented in Fig 2. The exact specification of this model based on Eqs (1)–(4) is given in S4 Text.

**Fig 3.**
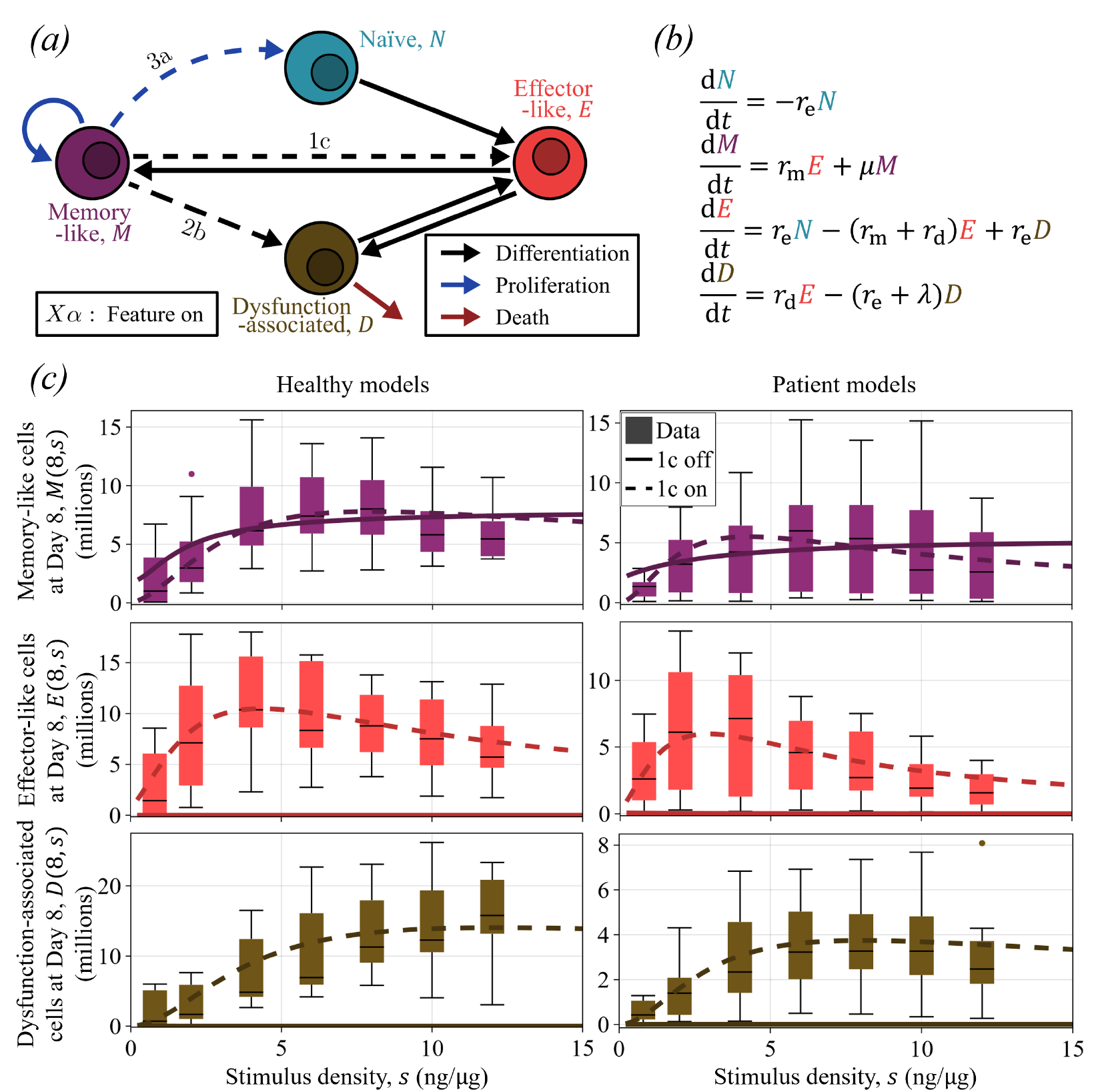
First case study. *(a)* Our interpretation of the pathway proposed by Wherry [57] (solid arrows), with alterations to the pathway (dashed arrows, conditions indicated with feature selections). *(b)* Ordinary differential equations that model the base pathway in *(a)* (solid arrows). *r*_m_, *r*_e_ and *r*_d_ are all dependent on stimulus density *s. (c)* Population predictions for memory-like, effector-like and dysfunction-associated cells at Day 8 with the base model (solid lines) and a modified model with feature 1c turned on (dashed lines), compared against Day 8 cell counts from the healthy (left) and the patient data sets (right) published by Zhang *et al*. [30].

After fitting the model in Fig 3b to data by minimising the healthy donor and cancer patient loss functions (green cross in Fig 2b), its predictions match neither the Day 8 cell counts for the healthy nor the patient data set across different stimulus densities, except for the memory-like cell compartment (solid lines in Fig 3c). This is likely due to the fact that memory-like cells are the only cells that self-renew in this model, and they do not transition to any other cell types to contribute to the expansion of their subpopulations. To address this, we considered introducing bidirectional differentiation (feature 1c) to this pathway by adding a transition from memory-like to effector-like states. With feature 1c turned on, re-fitting the model to data (green circle in Fig 2b) makes it match the healthy and patient data set more closely for the memory-like, effector-like and dysfunction-associated T cell compartments (dashed lines in Fig 3c), which supports the idea that memory-like cells may replenish the pools of other T cell phenotypes. We omit the naïve cell count predictions as they represent a negligible proportion of T cells expanded polyclonally.

As our second case study, Sun *et al*. [66] propose multiple T cell differentiation pathways under different contexts. They suggest that, during acute infections, naïve cells can differentiate to effector or memory precursor cells (which we assume to be memory-like for simplicity), which then both transition to memory-like cells after clonal contraction. In chronic infections and cancer, memory precursor cells are reported to progressively differentiate through dysfunctional states. We interpret these suggestions to form the base pathway (solid arrows) and mathematical model in Figs 4a and 4b, again under the assumption that dysfunction-associated cells approximate dysfunctional cells. The predictions made from this model (orange cross in Fig 2b, and solid lines in Fig 4c) may be sufficient to predict the Day 8 cell counts for memory-like and dysfunction-associated cells across different stimulus densities, but it predicts zero effector-like cells. Switching the direction of differentiation between memory-like and effector-like cells (turning feature 1a off) maintains similar predictions for memory-like and dysfunction-associated cell counts and also improves the prediction for the effector-like subpopulation (orange circle in Fig 2b, and dashed lines in Fig 4c).

**Fig 4.**
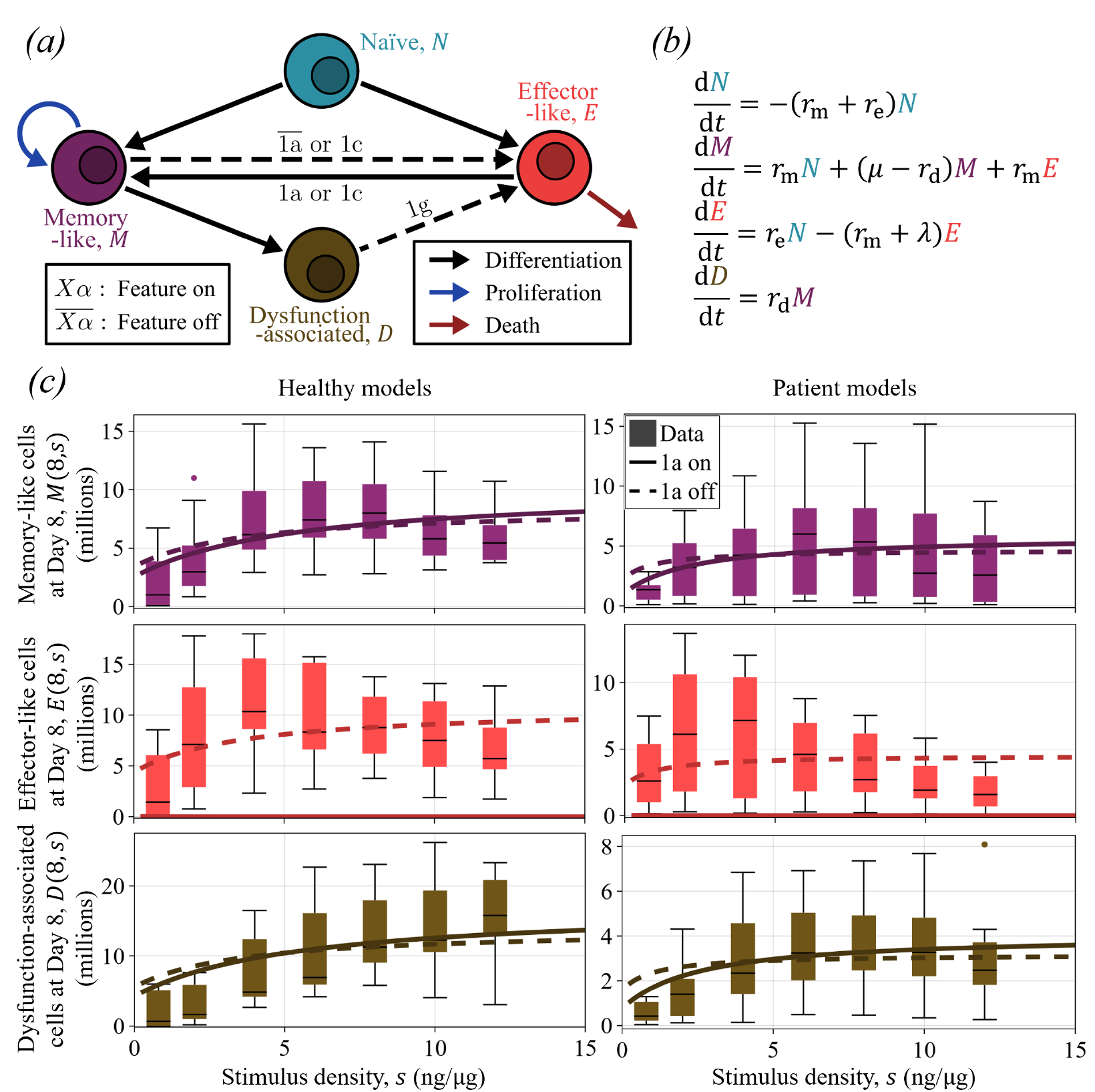
Second case study. *(a)* Our interpretation of the pathway proposed by Sun *et al*. [66] (solid arrows), with alterations to the pathway (dashed arrows, conditions indicated with feature selections). *(b)* Ordinary differential equations that model the base pathway in *(a)* (solid arrows). *r*_m_, *r*_e_ and *r*_d_ are all dependent on stimulus density *s. (c)* Population predictions for memory-like, effector-like and dysfunction-associated cells at Day 8 with the base model (solid lines) and a modified model with feature 1a turned off (dashed lines), compared against Day 8 cell counts from the healthy (left) and the patient data sets (right) published by Zhang *et al*. [30].

We consider multiple variations of the base pathways shown in Figs 3a and 4a by toggling only one other feature selection on or off (only those that substantially alter the model predictions). Table 3 summarises the normalised loss for each pathway and their ranking among all 7,840 models in terms of loss against each data set (lower loss corresponds to higher rank). For both case studies, introducing transitions from memory-like cells to another cell compartment, specifically bidirectional exchanges between memory-like and effector-like cells (feature 1c on), reduces model loss as it allows the expansion of other T cell subpopulations. Notably, the base model in the first case study (solid arrows in Fig 3a) ranks poorly at #7,482 among all models fitted to the patient data set, but upon introducing bidirectional differentiation (feature 1c on), this ranking improves to #14. Similarly for the second case study, the base model improves from #6,530 to #2 among the models fitted to the healthy data set when introducing bidirectional differentiation.

**Table 3.** Comparison between variations of differentiation pathways in the biological literature. Normalised minimal healthy donor and patient losses (Eq (9)) alongside their ranking among models fitted to each data set for variations of the base pathways in case study 1 (Fig 3) and case study 2 (Fig 4). The coloured markers correspond to models in Fig 2.

| Case study | Altered feature selection | Healthy data set |  | Patient data set |  |
| --- | --- | --- | --- | --- | --- |
| | | Normalised loss $\bar{\mathcal{L}}^h$ | Ranking | Normalised loss $\bar{\mathcal{L}}^p$ | Ranking |
| 1 | Base pathway (✕) | 3.737 | #7,480 | 3.466 | #7,482 |
|  | Feature 1c on (●) | 0.052 | #119 | 0.101 | #14 |
|  | Feature 2b on | 0.236 | #855 | 0.572 | #1,804 |
|  | Feature 3a on | 0.186 | #726 | 0.22 | #618 |
| 2 | Base pathway (✕) | 1.829 | #6,530 | 2.694 | #7,267 |
|  | Feature 1a off (●) | 0.426 | #2,535 | 0.673 | #2,392 |
|  | Feature 1c on | 0.035 | #2 | 0.149 | #373 |
|  | Feature 1g on | 0.166 | #690 | 0.515 | #1,393 |

### Differentiation features among the accepted models

The case studies indicate that small alterations in the structure of the differentiation pathway have the potential to significantly impact model predictions. Here we investigate common features among the models that most closely match both the healthy and patient data sets (those with low 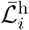 and 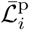 in Fig 2b). Fig 5a shows the average of the normalised healthy donor and patient losses 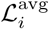 (Eq (10)). We see that the models with lowest average loss tend to be those whose predictions lie within one standard deviation about the mean of the data (blue), but some have predictions that lie outside one standard deviation (red).

**Fig 5.**
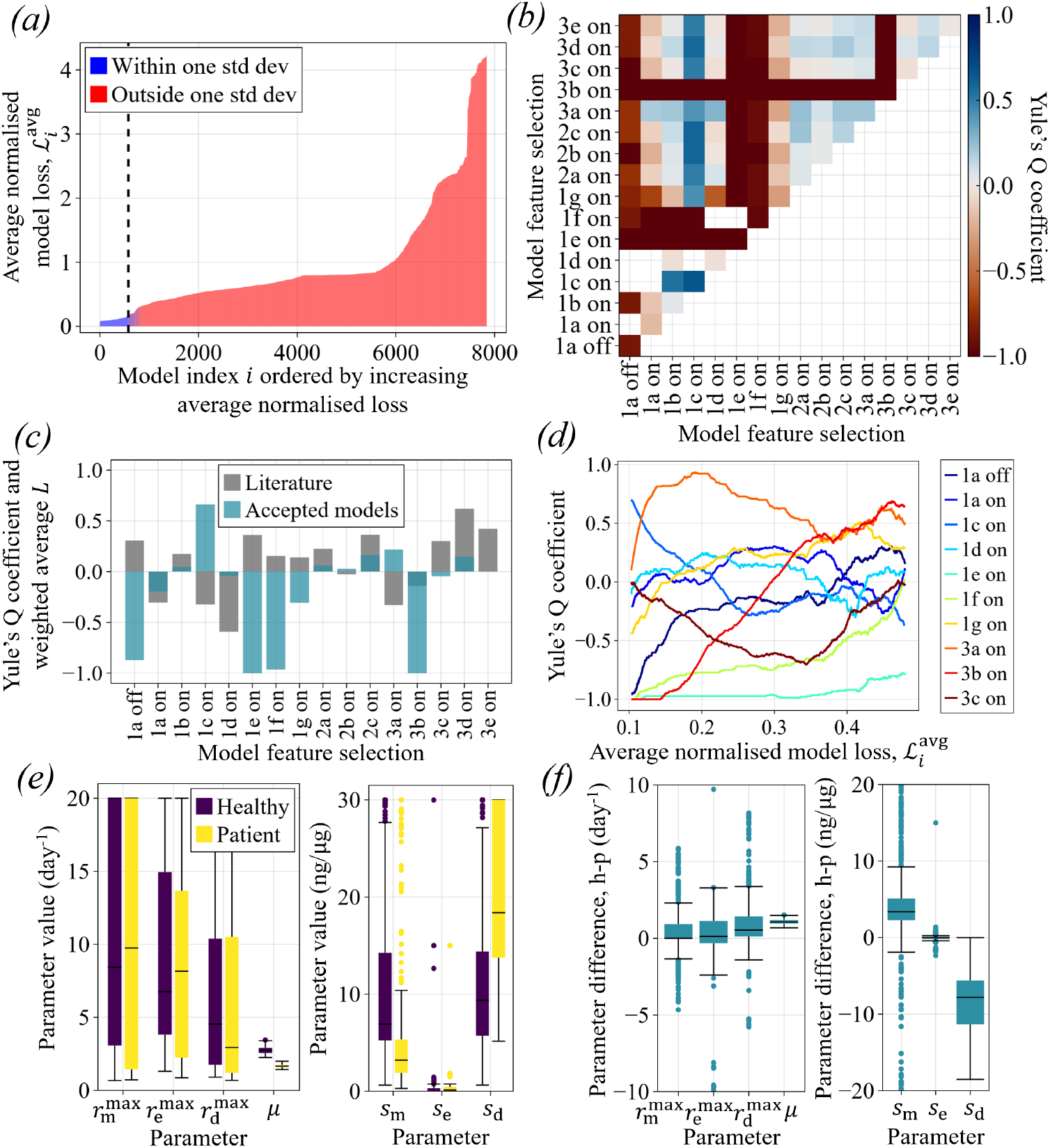
Feature selection frequency and parameter values among accepted models. *(a)* Average normalised model loss 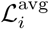 (Eq (10)) ordered by increasing loss (accepted 578 models are left of the vertical dashed line), coloured by models whose Day 8 predictions lie within (blue) or outside (red) one standard deviation about the mean of both data sets for all stimulus densities and all T cell counts except naïve cells. *(b)* Yule’s Q coefficient (Eq (11)) for pairs of feature selections in the accepted models. *(c)* Yule’s Q for each feature selection with the likeliness according to literature, computed as weighted average *L* (Eq (12)). *(d)* Yule’s Q for some feature selections in the 2,000 models with lowest average loss (vertical axis represents *Q* calculated over a 500 model window, horizontal axis is the average loss over the 500 models). *(e)* Parameter values and *(f)* differences in parameter values—healthy (h) minus patient (p) model parameters—among the accepted models (with some outliers omitted). 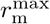 and *s*_m_ are the maximum rate and half-maximum stimulus density for differentiation to a memory-like state (similarly for parameters relating to effector and dysfunction-associated states, see Eqs (5)–(7)), and *µ* is the proliferation rate.

We conduct model selection to identify models that sufficiently approximate *ex vivo* T cell differentiation in the expansion experiments by Zhang *et al*. [30]. In ascending order of average loss 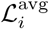, we sequentially check the predictions made by each model against both the healthy and patient data sets (with the corresponding fitted parameters for those data sets). We add models to our list of accepted models until 1% of them do not have predictions that fall within one standard deviation of the data mean for every relevant cell compartment, excluding naïve cells, and every stimulus density. This yields 578 models (those left of the vertical dashed line in Fig 5a), which we refer to as accepted models. All pathways ordered by average normalised model loss are shown in S3 Video, and the predictions made by the accepted models are shown in S1 Figure.

Fig 5b shows Yule’s Q coefficient (Eq (11)) for pairs of feature selections in the accepted 578 models, with diagonal entries representing individual feature selections. Entries without a listed value are either repeated pairs of feature selections (bottom right) or represent excluded feature combinations. Pairs with |*Q*| ≫ 0 have strong evidence for or against them in the context of short-term *ex vivo* expansion. Some individual feature selections with large |*Q*| seem to be dominant, in that models with these feature selections are likely to be among the accepted models regardless of most or all other selections. These dominant features include:

- differentiation from naïve to effector-like states with bidirectional exchanges between memory-like and effector-like compartments (feature 1c on)
- rate of memory-like cell differentiation increases with stimulus (feature 1e off)
- stimulus-dependence for differentiation to memory-like state and upregulation of inhibitory receptors (feature 1f off), but stimulus-independence for differentiation to effector-like state (*s*_e_ ≈ 0, see Fig 5e)
- no effector-like cell proliferation (feature 3b off).

Other feature selections appear in most of the accepted models but are less dominant, categorised by lower but non-negligible |*Q*| for most combinations with other features. These include:

- no downregulation of inhibitory receptors or differentiation from dysfunction-associated cells to effector-like states (feature 1g off)
- upregulation of inhibitory receptors among effector-like cells (feature 2c on)
- production of less differentiated daughter cells (assumed to be naïve-like) by proliferating cells (feature 3a on)

The accepted models do not strongly support a linear ordering of naïve, memory-like and effector-like compartments (feature 1a not on nor off). This is largely because models with bidirectional exchanges between memory and effector compartments dominate the accepted set of models, making it difficult to interpret feature 1a as on or off. Yule’s Q coefficient for pairs of feature selections in the top 578 models fitted to the healthy data set and the top 578 models fitted to the patient data set are given in S2 Figure, which shows very similar results to Fig 5b. This further supports the idea that the differentiation pathways that healthy and patient-derived T cells follow share similar structures.

Some feature selections have low |*Q*| for most combinations, which indicates that models may accurately describe the experimental data with or without these feature selections and we are unable to make conclusions regarding these selections. Consequently, our framework provides:

- no evidence for or against branching differentiation from naïve cells to memory-like and effector-like states (feature 1b), and no evidence for or against differentiation between memory-like and effector-like compartments (feature 1d)
- no evidence for or against the upregulation of inhibitory receptors on naïve (feature 2a) and memory-like (feature 2b) cells
- no evidence for or against naïve cell proliferation (feature 3c)
- no evidence for or against the death of effector-like (feature 3d) and dysfunction-associated cells (feature 3e).

We use these results to reconstruct a pathway diagram that represents the structure of our accepted models (Fig 6). This pathway shares similarities with the altered pathway from the first case study (bidirectional pathway in Fig 3a, and green circle in Fig 2b), which has the lowest average normalised loss out of all models.

**Fig 6.**
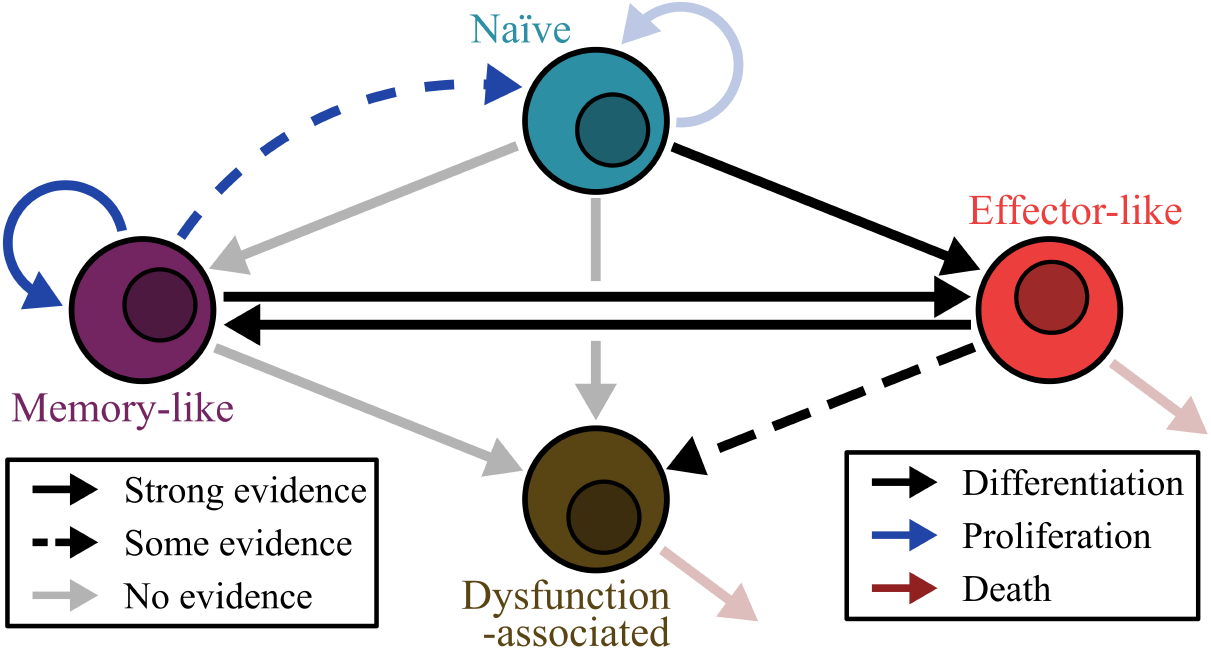
T cell differentiation pathway constructed from our model predictions. Hypothesis for the differentiation pathway connecting naïve, memory-like, effector-like and dysfunction-associated phenotypic compartments with transitions informed by the structures of the best-fitting models. Transitions, proliferation and death events related to features with relatively strong evidence (large Yule’s Q coefficient) are indicated by solid arrows, those with some evidence are indicated with dashed arrows, and those that we could not draw conclusions for are indicated with faded arrows. Features with evidence against them (1g and 3b) and those that do not alter pathway structure (1e and 1f) are not represented here. Note that the only hypothesised path leading to dysfunction is somewhat established (see Table 2).

Fig 5c shows Yule’s Q coefficient for individual feature selections (diagonal values in Fig 5b) alongside the likeliness of feature selections according to our interpretation of the literature *L* (Eq (12)). We see agreement between our findings and the literature (*Q* and *L* have the same sign) for around half of our features, and disagreement in the other half. Of the most significant features in our framework, only insignificant effector-like cell proliferation (feature 3b off) is supported by the literature on average. We also see some agreement that differentiation may not progress from naïve to effector-like to memory-like phenotypes (feature 1a not on), effector-like cells may upregulate inhibitory receptors (feature 2c on), and effector-like cells may die (feature 3d on) during short-term expansion. Although the literature seems to support these features on average, opinions on all features are relatively split in the literature (Table 2).

Fig 5d shows Yule’s Q coefficient for some feature selections (all features may be seen in S3 Figure) over samples of 500 models as the average model loss increases for the first 2,000 models (those that are moderately accurate when compared to the data). We see large variability in the frequency of some feature selections as the models generally worsen in their predictions, however, some remain relatively consistent. Although features 3a and 3b have relatively low |*Q*| for the 578 accepted models (Figs 5b and 5c), we see that they have large |*Q*| among models with average normalised losses between 0.2 and 0.4. Specifically, it is more likely for a model to be among moderately accurate (but not accepted) models if it includes asymmetric-like cell division (feature 3a on) or does not include naïve cell proliferation (feature 3c off).

Model predictions from the 1,000 models with the highest average loss 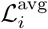 are shown in S4 Figure. These results alongside the case studies in Figs 3 and 4 indicate that, although some models are capable of capturing the population dynamics of some T cell subtypes, they do not explain the data for all subtypes. S5 Figure summarises the frequency of feature selections and parameter values among the 1,000 least accurate models. Some of the conclusions that may be drawn from these models are opposite to those in Fig 5, which is further evidence for those results. Specifically, Yule’s Q coefficient switches sign for features 1a on (supporting differentiation from naïve to memory-like to effector-like states), 1c (supporting bidirectional differentiation between memory-like and effector-like compartments), 2c (supporting the upregulation of inhibitory receptors in effector-like cells), and 3a (supporting asymmetric-like cell division).

### Differentiation kinetics among the accepted models

The rates of differentiation, dysfunction and proliferation associated with each model seem to vary depending on whether they were fitted to the healthy or patient data set. Although we expect that not all parameters are necessarily identifiable in most of our models, we may gain a general understanding of T cell differentiation kinetics across the accepted 578 models (Fig 5e). We estimate the T cell proliferation rate to range from around 1.5 to 3.5 day^−1^, which is supported by other *ex vivo* experiments [97–99]. We find that the maximum rates of differentiation are typically large, which is expected as we see rapid expansion of each T cell subtype in experiments [30]. Interestingly, both healthy and patient-derived T cells seem to differentiate to effector-like states at the maximal rate independent of T cell receptor stimulation since *s*_e_ ≈ 0 for most models.

We also consider the differences in parameter values between models fitted to the healthy and patient data sets (healthy minus patient model parameters, Fig 5f). In our models, T cells derived from healthy donors seem to proliferate more rapidly than patient-derived T cells. Additionally, the model indicates that healthy cells may require more stimuli to differentiate to memory-like states, and less stimuli to begin upregulating inhibitory receptors than patient cells. The maximum rates of differentiation and upregulation of inhibitory receptors seem consistent across healthy and patient-derived T cells. Among the worst 1,000 models (S5 Figure), we see that *s*_e_ is non-zero, the proliferation rate is approximately the same for healthy and patient T cells, and that the sensitivity to stimulus for differentiation does not differ significantly between healthy and patient cells, which supports our conclusions from Fig 5f.

### Biologically informed constraint of feature selection

Although our methods reveal information about some features of the T cell differentiation pathway, there is still uncertainty in our results and we are unable to draw conclusions regarding some features due to a lack of evidence. If it is the case that our conclusions are incorrect regarding one of the dominant feature selections (those with large |*Q*| in Fig 5b), examining the features and kinetics of all accepted models that fix this selection may reveal more information about features that are sub-dominant. Specifically, one may reason that the differentiation pathway for short-term *ex vivo* expansion might not involve bidirectional differentiation (contrary to our findings in Fig 5), as effector-like cells are often thought to only transition to memory-like states after clonal contraction [27, 57, 66, 69]. We investigate the impacts of this assumption here.

Fig 7 summarises the features present among the 204 accepted pathways that do not include bidirectional differentiation between memory-like and effector-like compartments (feature 1c off). Fig 7a indicates that some hypotheses regarding feature likeliness change under this constraint, namely:

- some evidence that differentiation might branch from naïve cells to memory-like and effector-like cells (feature 1b on)
- some evidence that differentiation might either progress from naïve to effector-like to memory-like states (feature 1a on), or that there is no differentiation between memory-like and effector-like compartments (feature 1d on with feature 1b)
- stronger evidence that naïve and effector-like cells might upregulate inhibitory receptors (features 2a and 2c on) and that memory-like cells might not (feature 2b)
- stronger evidence that proliferating cells might produce less differentiated daughter cells, which we assume to be naïve-like (feature 3a on).

**Fig 7.**
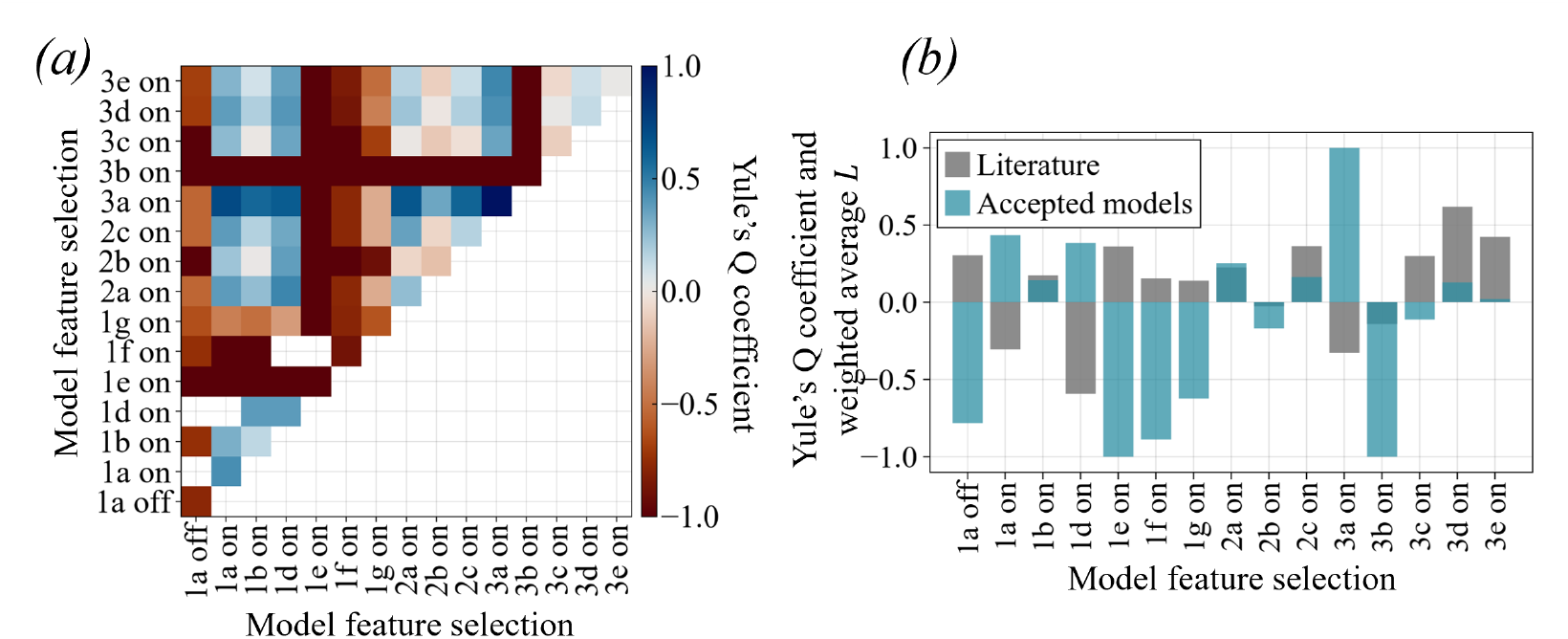
Frequency of feature selections among the accepted models without bidirectional differentiation. *(a)* Yule’s Q coefficient (Eq (11)) for pairs of pathway feature selections in the accepted models with feature 1c off (compared against all models with feature 1c off). *(b)* Yule’s Q coefficient for each feature selection in the accepted models with feature 1c off, plotted alongside the likeliness according to literature, computed as weighted average *L* (Eq (12)).

We do not see much change in parameter values upon assuming that the pathway does not involve bidirectional differentiation (results not shown). Although some conclusions change regarding the likeliness of feature selections, the literature still agrees with around half of these findings on average (Fig 7b).

## Discussion

In this work, we compare 7,840 literature-informed mathematical models of T cell differentiation to experimental data describing the *ex vivo* expansion of healthy and patient-derived T cells. Despite non-trivial differences between healthy and patient T cell counts, we find a strong correlation between the healthy and patient losses. This suggests that the immune dynamics within cancer patients prompt modifications to the regulatory dynamics of T cell differentiation and dysfunction rather than pathway structure. Biological studies have also hypothesised that regulatory T cells follow similar developmental pathways regardless of their origin or experience [100, 101], and that dysfunction arises from inherent properties of T cell responses that are present during both acute and chronic stimulation [52].

Our model suggests that healthy T cells likely proliferate more rapidly, and might require more stimulation for differentiation to memory-like states and less for the upregulation of inhibitory receptors than cancer patient T cells. The latter may be a controversial result as antigen-experienced cells are typically thought to be more sensitive to antigenic stimulation than inexperienced cells [102], but a study by Mehlhop-Williams and Bevan suggests that antigen-experienced cells may be less sensitive for initiating proliferation [103]. Although this does not directly indicate a relationship between T cell origin and dysfunction, it suggests that there may be more to learn regarding stimulation sensitivity for T cell differentiation. Developing our understanding of the differences between healthy and patient-derived cells is an important step towards designing the expansion of tumour-infiltrating lymphocytes such that cancer-specific T cells are selectively enriched.

By observing the structures of models that sufficiently explain experimental observations, our framework can provide biologically testable hypotheses regarding short-term T cell differentiation in an *ex vivo* setting under interleukin-2 supplementation and polyclonal stimulation with anti-CD3 and anti-CD28. Significantly, models containing bidirectional exchanges between memory-like and effector-like phenotypic compartments are enriched among the best-fitting models under the study’s assumptions. This suggests that T cells may differentiate along a route that is more dynamic and plastic than a simple linear trajectory. The bidirectional trajectory allows for memory-like cells to support the rapid expansion of other T cell phenotypes as seen in experiments. Bidirectional differentiation is typically only discussed in the context of long-term *in vivo* immune responses [2, 4, 5], however, we would not expect clonal contraction and memory formation at Day 8 in *ex vivo* experiments. To test this hypothesis, longitudinal analyses of T cell phenotypes may help to reveal how memory-like and effector-like cells differentiate over time during *ex vivo* expansion.

We find that our computational models are more likely to explain experimental observations if effector-like cells do not proliferate and if they upregulate inhibitory receptors during *ex vivo* expansion, which is supported in the literature [8, 57, 59, 81]. Our framework indicates evidence of stimulus-dependence for differentiation to memory-like states and upregulation of inhibitory receptors but stimulus-independence for differentiation to effector-like states. This may be consistent with evidence that, after sufficient initial T cell receptor stimulation, activated cells can become committed to subsequent proliferation and differentiation without continuous antigenic stimulation [5, 77]. Models not containing back-differentiation from dysfunction-associated to effector-like cells are enriched among the accepted models, which raises the hypothesis that T cells co-expressing PD-1 and TIM-3 may not downregulate these inhibitory receptors (or that the back-differentiation of dysfunction-associated cells is insignificant compared to differentiation from other cell states into effector-like states). Our findings also suggest that activated cells may be maintaining a pool of less differentiated cells, which could be explained by the re-expression of markers associated with less differentiated cells, or compensation for a missing subset-specific proliferation term in the model. Some studies suggest that asymmetric cell division is prominent in environments with high stimulus density [78, 85], which could also explain this result. Further biological testing using lineage tracing may help to identify the cause of this observation.

Upon assuming that the T cell differentiation pathway does not involve bidirectional exchanges, we hypothesise that the differentiation pathway has a branching structure. Although this prediction assumes prior knowledge of one feature in the differentiation pathway, it indicates that we are able to reduce uncertainty in our results by using information gained by future biological experimentation. Specifically, we suggest that biological research focused on understanding bidirectional phenotype switching, differentiation kinetics (specifically the dependence of T cells on polyclonal stimuli) and effector-like cell proliferation could help to improve the predictions of our mathematical framework. Longitudinal sampling, lineage tracing, proliferation tracking, and functional measurements could discriminate among the leading models.

## Conclusion

In this work, we systematically translate a broad and often conflicting biological literature into a large set of candidate T cell differentiation pathways which we evaluate within a common framework rather than testing only one or two predefined models. By considering multiple plausible pathways, the study captures uncertainty in our current understanding and identifies mechanisms that are consistently compatible with the experimental data by Zhang *et al*. [30] and with questions that remain unresolved. Our computational model integrates differentiation-associated phenotypes, inhibitory receptor expression, stimulation density, and T-cell source, allowing these processes to be examined together rather than in isolation. Importantly, the framework generates experimentally testable hypotheses and identifies the measurements most likely to distinguish among competing biological explanations.

We present three main findings. First, the same pathway structures tend to explain the endpoint phenotypes of both healthy and patient-derived T cells, while their estimated kinetic parameters differ. This raises the testable hypothesis that donor health status and treatment history primarily alter the rates and stimulus responsiveness of phenotypic changes, rather than the repertoire of possible state transitions.

Second, our analysis of feature frequency in the best-fitting models identifies two biological processes that may be especially important during *ex vivo* expansion. These include transitions between memory-like and effector-like marker-defined states, suggesting that the balance between these populations may be dynamic rather than unidirectional. The accepted models also preferentially place the acquisition of PD-1 and TIM-3 within the effector-like compartment, raising the hypothesis that effector differentiation may increase the likelihood of developing an inhibitory-receptor-positive phenotype. Third, the finding that explicit effector-like proliferation is not required in the best-fitting models suggests that the expansion of effector-like populations may be maintained by other compartments or transitions. These predictions are experimentally testable using longitudinal phenotyping, lineage tracing, and functional assays. Our framework identifies which biological uncertainties most strongly affect interpretation of the expansion data (specifically memory and effector state switching and subset-specific proliferation), which highlights this study’s value as a tool for prioritizing future experiments to reduce uncertainty.

Our study relies on limited experimental data and a number of assumptions, and therefore has some limitations. Our hypotheses regarding T cell differentiation dynamics are likely to be affected by the lack of time-series information in the data sets to which we fitted our models. Similarly, the data does not contain counts for dead cells as they were gated out in a preliminary stage of flow cytometry, which may have caused uncertainty regarding cell death in our model. With measurements primarily at baseline and Day 8, changes in the proportions of marker-defined populations cannot distinguish true differentiation from differential proliferation, survival, or transient changes in marker expression. For example, an increase in memory-like cells could reflect differentiation into that state, preferential expansion of pre-existing memory-like cells, or greater survival. Similarly, the data cannot directly demonstrate bidirectional transitions between memory-like and effector-like cells without longitudinal or lineage-tracing measurements. Additionally, we only consider modelling discrete naïve, memory-like, effector-like and dysfunction-associated cell compartments, and we grouped cells expressing CD4 and CD8. These limitations may explain the disagreements between our findings and the biological literature, and the inability of our framework to provide evidence for some aspects of T cell differentiation.

Future work could include expanding our mathematical model with more distinct T cell phenotypes including a broader range of dysfunction-associated cell states, and fitting these models to experimental data with more extensive time-series information. Our models may be used to predict experimental alterations that are likely to improve the expansion of memory-like cells and reduce the expression of inhibitory receptors among expanded populations while accounting for uncertainty in the structure of the differentiation pathway. Since we found that the same mathematical models explain data for T cells of varying origins, these analyses may translate to the expansion of T cells originating from individuals with cancer types not represented in the data. Our framework may be extended to propose hypotheses regarding T cell differentiation and inhibitory receptor regulation during long-term *in vivo* immune challenges, with antigen-specific stimulation, or in the presence of other cytokines. Its application to healthy and patient-derived cells also gives this work direct translational relevance, as it could ultimately help optimize stimulation conditions and culture duration to preserve desirable T cell states during patient-specific CAR T cell manufacturing. The hypotheses generated using our method have the potential to significantly expand our understanding of T cell differentiation which may be exploited to improve the efficacy of cancer immunotherapies.

## Supporting information

**S1 Text. Literature review**. Justifications for the citations in Table 2 relating to the estimated likeliness of pathway features being on during short-term *ex vivo* T cell differentiation.

**S2 Text. Calculation of cell counts from data**. Assumptions and calculations used to determine cell counts for naïve, memory-like, effector-like and dysfunction-associated cells from the data published by Zhang *et al*. [30].

**S3 Text. Excluded pathway feature combinations**. List and justifications of the combinations of pathway features that were excluded from this study.

**S4 Text. Mathematical model formulation**. Elaborated description for the formulation of the mathematical model in Eqs (1)–(4), and the derivation of the models in Figs 3b and 4b.

**S5 Text. Details for solving the mathematical models and parameter estimation**. Calculations used to solve the ordinary differential equation system in Eqs (1)–(4) at Day 8 and derive the gradient of the loss function in Eq (8) to optimise computational efficiency during model fitting.

**S1 Video. All pathways ordered by increasing healthy donor loss**. Animation showing the structure of the differentiation pathways corresponding to models in order of increasing healthy donor loss 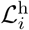 (Eq (8), most to least accurate models). Dysfunction-associated cells are denoted as dysfunctional for brevity.

**S2 Video. All pathways ordered by increasing patient loss**. Animation showing the structure of the differentiation pathways corresponding to models in order of increasing patient loss 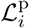 (most to least accurate models). Dysfunction-associated cells are denoted as dysfunctional for brevity.

**S3 Video. All pathways ordered by increasing average normalised model loss**. Animation showing the structure of the differentiation pathways corresponding to models in order of increasing average normalised model loss 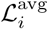 (Eq (10), most to least accurate models). Dysfunction-associated cells are denoted as dysfunctional for brevity.

**S1 Figure. T cell subpopulation predictions from the accepted models**. Predictions of Day 8 cell counts for *(a)* naïve, *(b)* memory-like, *(c)* effector-like and *(d)* dysfunction-associated cells as stimulus density varies using the accepted 578 models (those with the lowest average model loss 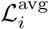) compared to cell counts computed from the healthy (left) and patient (right) data sets published by Zhang *et al*. [30].

**S2 Figure. Frequency of pairs of features in models fitted to the healthy and patient data sets**. Yule’s Q coefficient (Eq (11)) for pairs of pathway feature selections in the 578 models with least healthy donor loss 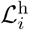 (left) and patient loss 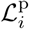 (right).

**S3 Figure. Frequency of features across the 2**,**000 most accurate models**. Yule’s Q coefficient (Eq (11)) for each feature selection in the 2,000 models with lowest average loss 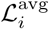 (vertical axis represents Yule’s Q coefficient calculated over a 500 model window, horizontal axis is the average loss over the 500 models).

**S4 Figure. T cell subpopulation predictions from the least accurate models**. Predictions of Day 8 cell counts for *(a)* naïve, *(b)* memory-like, *(c)* effector-like and *(d)* dysfunction-associated cells as stimulus density varies using the 1,000 models with the highest average model loss 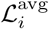 (Eq (10)) compared to cell counts computed from the healthy (left) and patient (right) data sets published by Zhang *et al*. [30].

**S5 Figure. Feature frequency and parameter values among the least accurate models**. *(a)*

Average normalised model loss 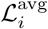 (Eq (10)) ordered by increasing loss, with the last 1,000 models indicated by the vertical dashed line, coloured by models whose Day 8 predictions lie within (blue) or outside (red) one standard deviation about the mean of both data sets for all stimulus densities and all T cell counts except naïve cells. *(b)* Yule’s Q coefficient (Eq (11)) for pairs of pathway feature selections in the 1,000 models with the highest average loss 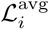. *(c)* Yule’s Q coefficient for each feature selection in the worst models plotted alongside the likeliness according to literature, computed as weighted average *L* (Eq (12)).

*(d)* Parameter values and *(e)* differences in parameter values—healthy (h) minus patient (p) model parameters—among the 1,000 least accurate models.

## Acknowledgments

P.R.B. and A.L.J. acknowledge support from the Australian Research Council (ARC) Discovery Project (DP) DP230100025. A.L.J. acknowledges support from the ARC Discovery Early Career Researcher Award (DECRA) DE240100650. M.S.L. acknowledges support from the QUT Centre for Data Science.

