## Supplementary figures and images for "A computational investigation of pathways for *ex vivo* differentiation and dysfunction of healthy and patient T cells"

### Supplementary Figure 1

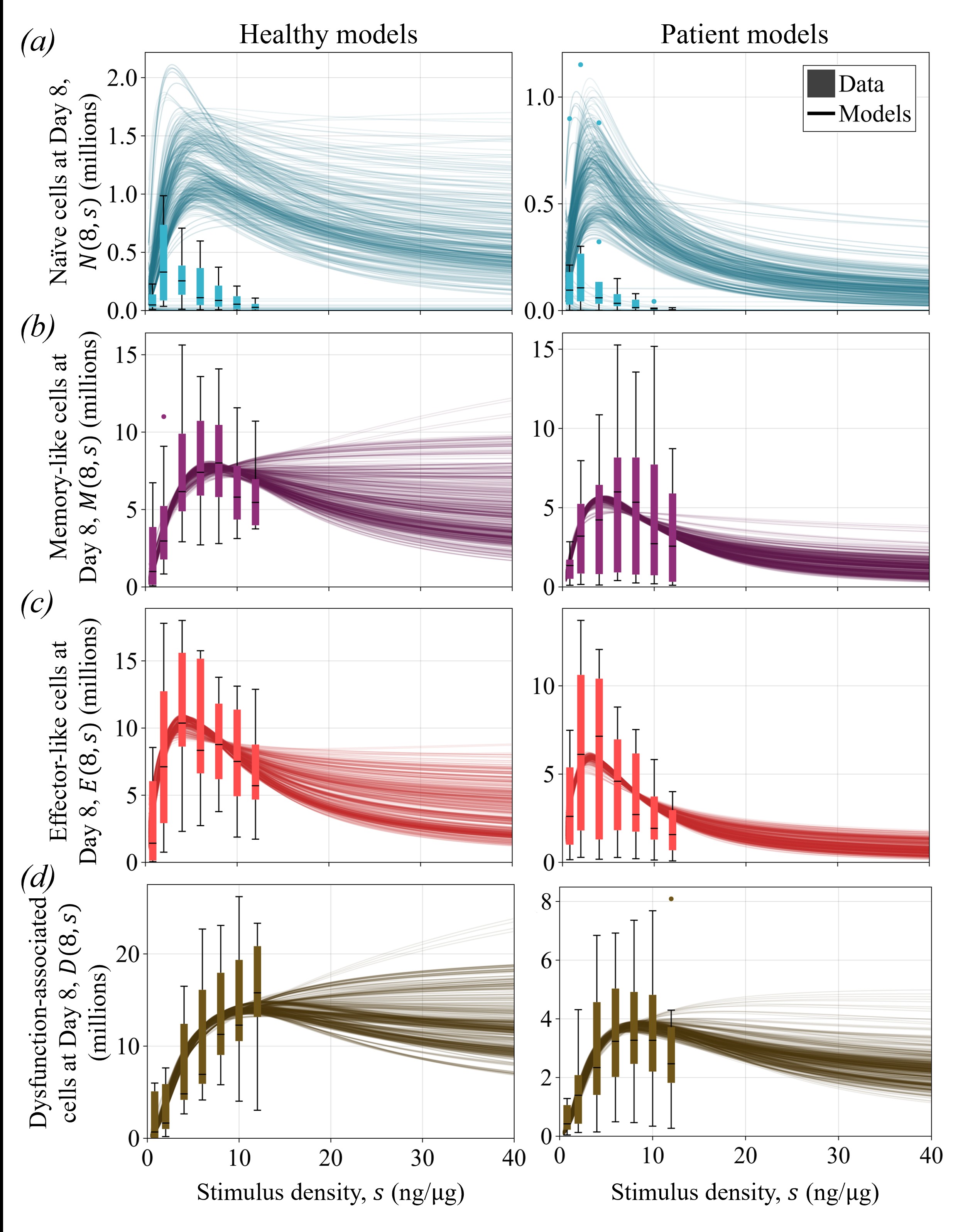

### Supplementary Figure 2

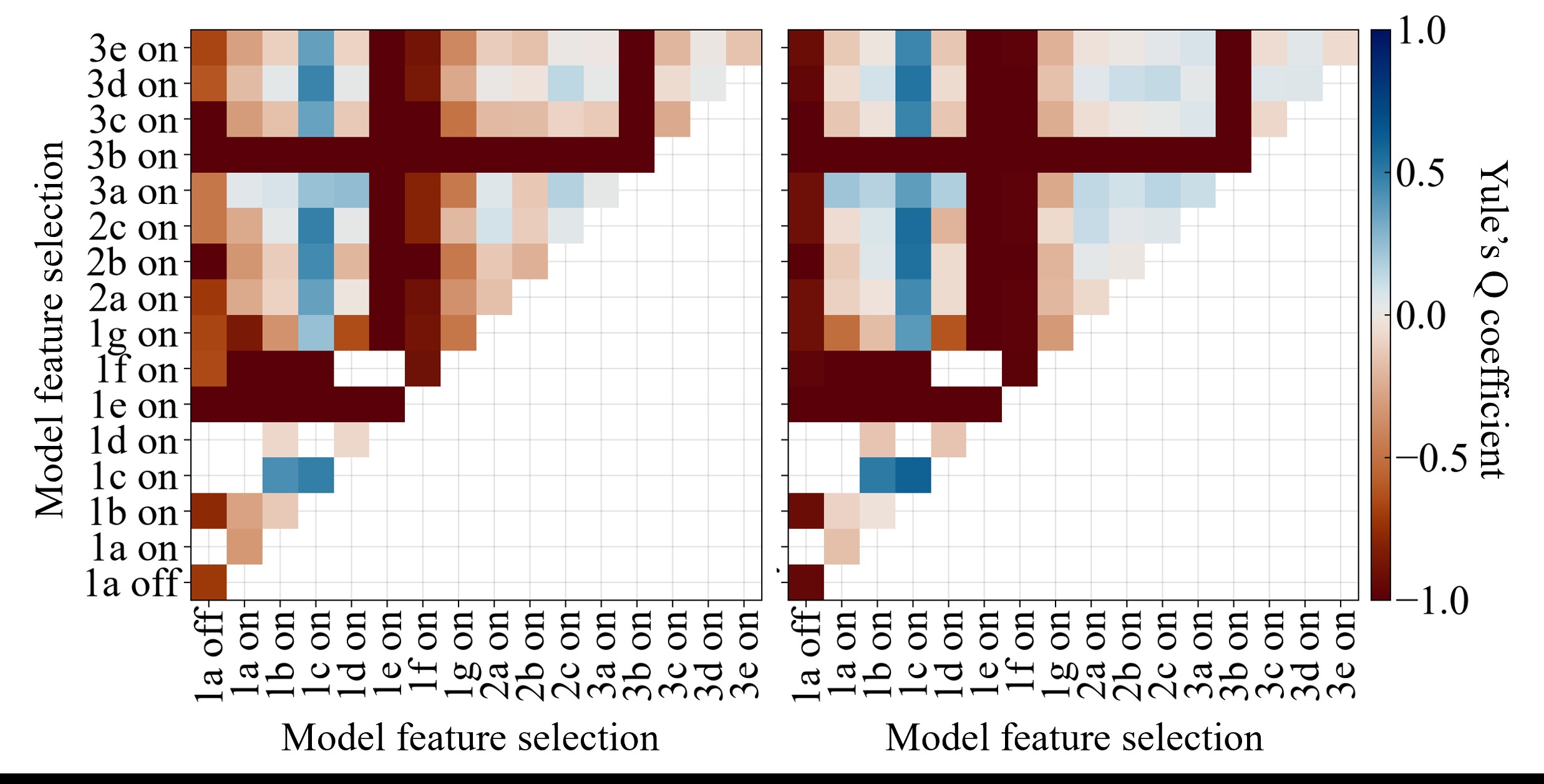

### Supplementary Figure 3

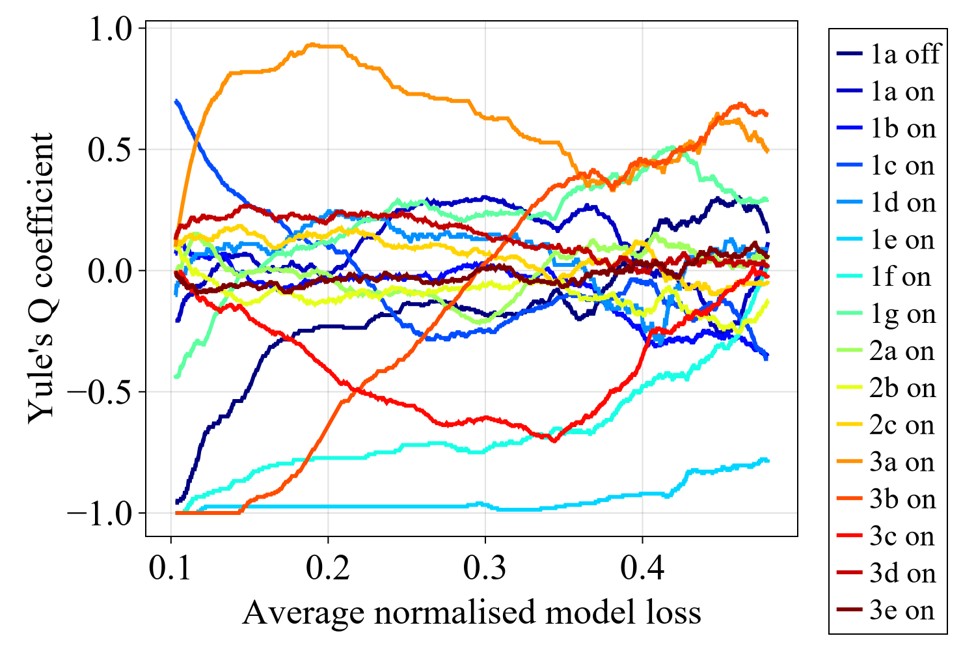

### Supplementary Figure 4

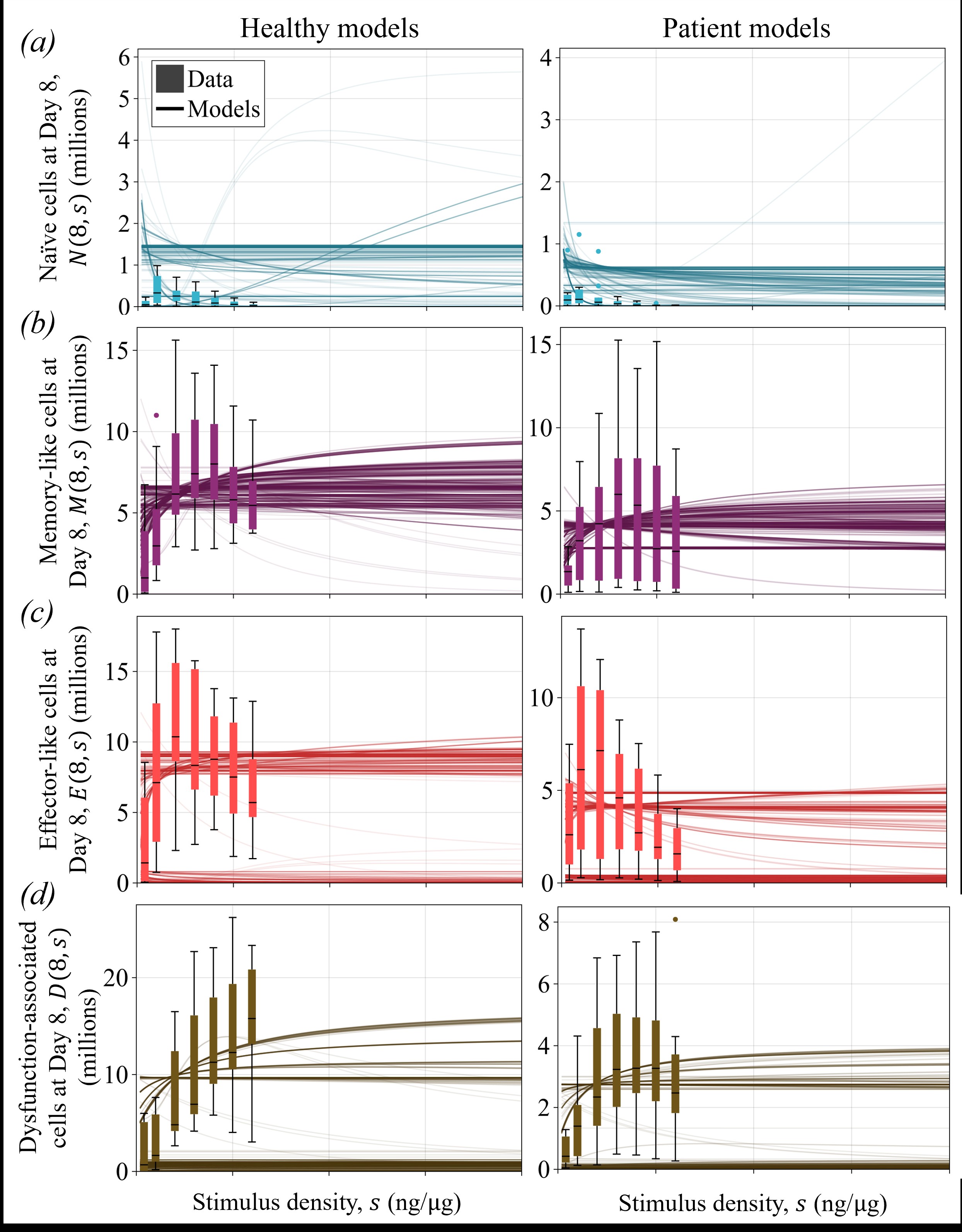

### Supplementary Figure 5

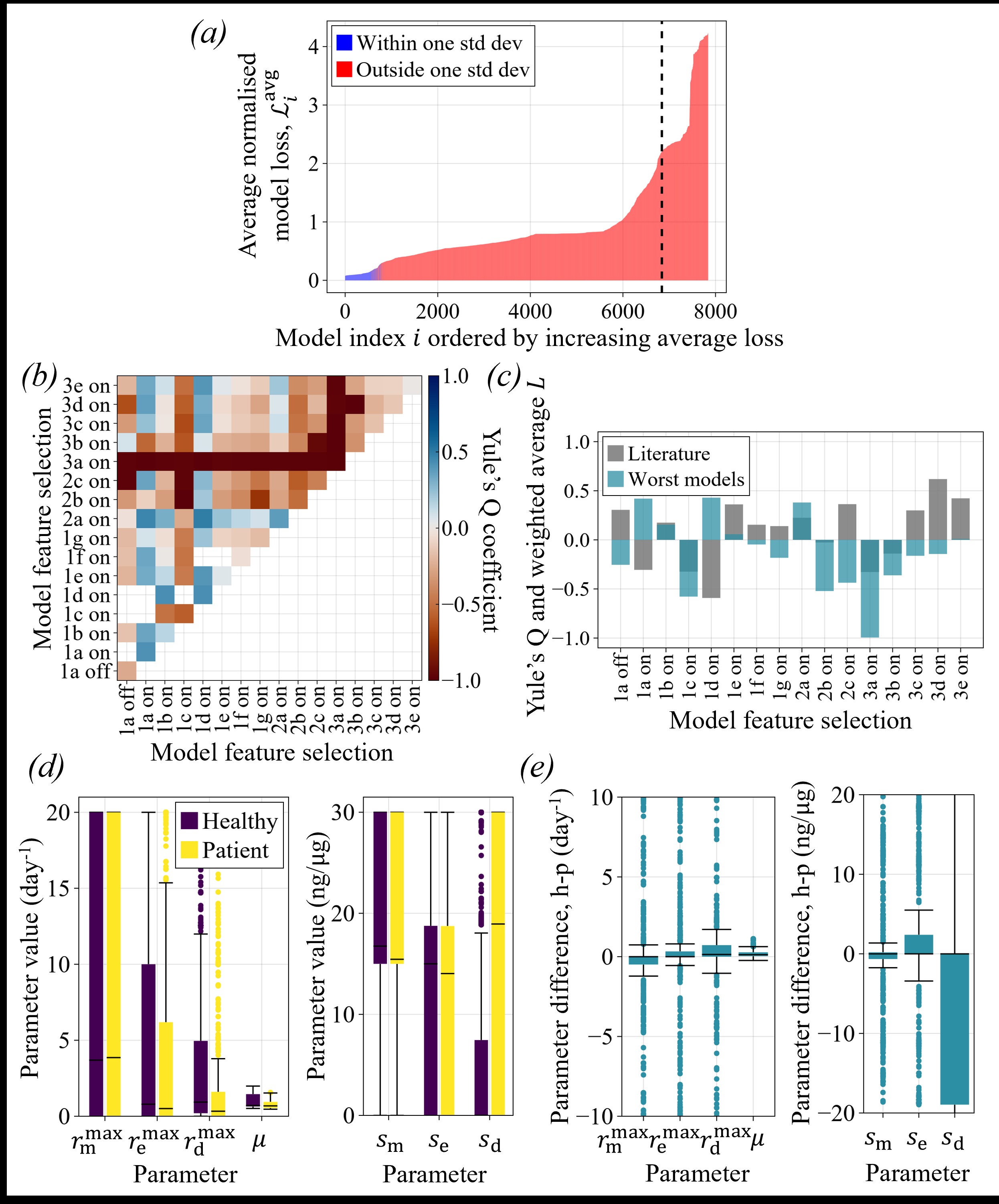
