## Supplementary Text 1 for "A computational investigation of pathways for *ex vivo* differentiation and dysfunction of healthy and patient T cells"

ORCIDs: MSL, 0009-0009-6633-2044; YB, 0000-0001-9901-7782; PRB, 0000-0003-3962-5393; ALJ, 0000-0001-9103-7092

October 19, 2026

### S1 Text. Literature review

The following lists detail our interpretations of each article cited in Table 2 of the main text relating to each relevant pathway feature in the context of short-term *ex vivo* expansion. The citations are ordered and numbered as they appear in the main text. If an article does not discuss or imply information about a feature, then we do not mention it here. For the purpose of deciding feature likeliness, we assume that the T cell differentiation pathway has the same structure during acute and chronic stimulation, for T cells expressing CD4 and CD8, and for T cell responses *in vivo* and *ex vivo*. For brevity, we refer to cells with memory-like and effector-like phenotypes (as defined in Table 1 of the main text) as memory and effector cells respectively. We assume that pathway features related to exhaustion, or dysfunction, also apply for our dysfunction-associated cell compartment (relating to the regulation of inhibitory receptors). Note that we did not interpret any articles to find feature 1e unlikely since, whenever articles mention stimulus dependence for differentiation, they typically indicate support for biological processes similar to that described by feature 1e. Similarly for feature 3d, whenever cell death is discussed, effector-like cells are indicated to be susceptible to cell death.

Henning *et al.* (2018) [2] propose multiple potential pathways for differentiation and suggest where dysfunction may occur in one of the pathways:

- They suggest the possibility of a bidirectional pathway in which memory cells only differentiate to effector states after a secondary immune challenge (which may not be the case during *ex vivo* expansion), and also a linear pathway in which greater stimulation results in more effector cells. This indicates that feature 1d is unlikely, but they do not indicate which pathway structure is more likely, so we assume that features 1a, 1c and 1e are undecided/neutral.
- They do not suggest a branching differentiation pathway, so we interpret this to implicitly mean that feature 1b is unlikely.
- Both models express conflicting opinions regarding antigen dependence for differentiation, so we assume that feature 1f is neutral.
- The pathway for dysfunction shows that early dysfunctional cells may back-differentiate to memory cells, which somewhat supports feature 1g (back-differentiation, but not necessarily to an effector-like state).
- They state that effector cells become dysfunctional and their diagram suggests that memory cells may also become dysfunctional, but they do not suggest that naïve cells do so, which implies that feature 2a is unlikely and that feature 2b is likely, and supports that feature 2c is likely.
- They propose the possibility of asymmetric cell division to explain heterogeneity in expanded cell pools, but also depict self-renewal (symmetric proliferation) in pathway diagrams, so we assume that feature 3a is neutral.
- The pathways depict memory cell proliferation, but not naïve or effector cell proliferation, which implies that features 3b and 3c are unlikely.
- They state that effector cells die which supports feature 3d, but they do not show dysfunctional cell death and imply that these cells only die after back-differentiating, which implies that feature 3e is unlikely.

Reiser and Banerjee (2016) [3] present a pathway for T cell differentiation and discuss dysfunction in the context of acute infection and antitumour responses:

- The pathway indicates that effector cells differentiate to memory cells after antigen clearance, which does not necessarily occur during *ex vivo* expansion, so we assume that feature 1a is implicitly likely and feature 1d is implicitly unlikely.
- They state that naïve cells directly differentiate to effector cells, which suggests that feature 1b is unlikely.
- They state that memory cells differentiate back into effector cells during antigen rechallenge, which somewhat (or implicitly) supports feature 1c.
- Since memory cells are stated to arise only after antigen is cleared, this suggests that effector cells require little antigen to differentiate to memory cells, so feature 1e is implicitly likely.
- They state that T cell exhaustion can be reversed, which supports feature 1g.
- They indicate that effector cells make up the majority of the cell pool that becomes dysfunctional, which implies that feature 2c is likely.
- They report that memory cells self-renew (which we assume to indicate symmetric proliferation),

so we assume that feature 3a is implicitly unlikely. They do not comment on the proliferation of other cell subtypes
- They state that effector cells die after antigen clearance (which does not necessarily occur during *ex vivo* expansion) and that exhausted cells eventually die, which somewhat (or implicitly) supports feature 3d and supports feature 3e.

Restifo and Gattinoni (2013) [4] propose two potential pathways for differentiation:
- Both pathways indicate branching differentiation and that memory cells differentiate to effector cells, which supports feature 1b and suggests that features 1a and 1d are unlikely.
- One potential pathway suggests bidirectional differentiation and the other does not, but they do not indicate which one is more likely, so we assume that feature 1c is neutral.
- The pathways indicate that memory cells self-renew (proliferate symmetrically) and that naïve and effector cells do so at a lower rate, which indicates that feature 3a is unlikely and somewhat supports that features 3b and 3c are likely.
- In one pathway, the effector cell state is terminal and these cells die, but in the other pathway, not all effector cells die, so we assume that feature 3d is implicitly likely.

Kedzierska and Koutsakos (2020) [5] propose multiple differentiation pathways:
- They do not suggest that one pathway is more valid than another, so we interpret this to mean that they have neutral opinions for features 1a, 1b, 1c, 1d and 1e (linear differentiation from memory to effector cells with more memory cells under less T cell receptor stimulation, bidirectional differentiation, and branching transitions from naïve cells to memory and effector cells without differentiation between memory and effector cells).
- Each pathway suggests that only the first differentiation event requires antigenic stimulation, which supports feature 1f.
- They suggest an asymmetric division model and do not suggest symmetric division in other models, so we assume that this implies support for feature 3a.
- Effector cells are said to have limited proliferative potential, which suggests that feature 3b is somewhat (or implicitly) unlikely.

Buchholz *et al.* (2016) [6] do not explicitly present a differentiation pathway, but they discuss ideas surrounding differentiation:
- They support multiple potential ideas, such as that T cells progressively differentiate from memory to effector cells with constant antigenic stimulation (relates to features 1a, 1d and 1f), and also an idea similar to branching differentiation in which differentiation leads to memory cells under less stimulation (features 1b and 1e), but since none are implied to be most likely, we consider these features to be neutral.
- They suggest that asymmetric naïve cell proliferation may be driving diversity in expanded cell populations, which implies support for features 3a and 3c.

Chen *et al.* (2026) [7] present pathways for differentiation during acute stimulation and dysfunction during chronic stimulation:
- They present a branching pathway without differentiation between memory and effector cells under acute antigenic stimulation, which supports features 1b and 1d.
- They do not suggest a bidirectional differentiation pathway, so we interpret this to implicitly mean that feature 1c is unlikely.
- Transitory dysfunctional cells may differentiate into dysfunctional cells with effector-like phenotypes, which implies support for feature 1g.
- Only naïve cells are shown to become early dysfunctional cells (memory and effector cells are not shown to do this), which implies support for feature 2a, and implies that features 2b and 2c are unlikely.
- Early dysfunctional cells are shown to self-renew (symmetrically proliferate) which we assume to also hold for memory-like cells, which suggests that feature 3a is implicitly unlikely.

Blank *et al.* (2019) [8] extensively discuss T cell dysfunction and present a pathway involving differentiation and dysfunction:

- They present a pathway which involves both branching differentiation and memory to effector cell differentiation under acute stimulation, which supports feature 1b, and suggests that features 1a and 1d are unlikely.
- They do not suggest a bidirectional differentiation pathway, so we interpret this to implicitly mean that feature 1c is unlikely.
- They suggest that it is unknown whether there are conversions between early dysfunctional cells and effector cells or not, which suggests that features 1g and 2c are undecided/neutral.
- The pathway indicates that naïve cells become dysfunctional under chronic stimulation but memory cells do not, which supports feature 2a and implicitly suggests that feature 2b is unlikely.
- The pathway depicts self-renewal (symmetric proliferation) of early dysfunctional cells which we assume to also hold for memory-like cells, which suggests that feature 3a is implicitly unlikely.
- They suggest that dysfunctional cells are less prone to cell death as dysfunction prevents differentiation into an effector-like state, which implies that effector cells die (feature 3d is likely) and that dysfunctional cell death is less significant (feature 3e is unlikely).

Ahmed *et al.* (2023) [22] present a pathway for T cell differentiation and dysfunction:

- They state that effector cells differentiate to memory cells after antigen clearance, which does not necessarily occur during *ex vivo* expansion, so we assume that feature 1a is implicitly likely and feature 1d is implicitly unlikely.
- The pathway shows branching differentiation from mature naïve T cells (after antigenic stimulation), which supports feature 1b.
- They suggest a pathway that does not involve bidirectional differentiation, so we interpret this to mean that feature 1c is implicitly unlikely.
- Since memory cells are stated to arise only after antigen is cleared, this suggests that effector cells require little antigen to differentiate to memory cells, so feature 1e is implicitly likely.
- They discuss the possibility of reversing T cell dysfunction, which supports feature 1g.
- The pathway indicates that effector cells become dysfunctional but it does not show pathways from naïve or memory cells to dysfunctional cells, so we assume this to mean that features 2a and 2b are implicitly unlikely and feature 2c is likely.
- They state that memory cells self-renew, which we assume implies that proliferation is symmetric and so feature 3a is implicitly unlikely.
- They report that effector cells die after antigen clearance (which does not necessarily occur during *ex vivo* expansion) and that exhausted cells are ready to die, so features 3d and 3e are implicitly likely.

Kishton *et al.* (2017) [23] propose a pathway for T cell differentiation:

- They propose a linear pathway in which memory cells differentiate to effector cells, which suggests that features 1a and 1d are unlikely.
- They do not suggest a branching or bidirectional differentiation pathway, so we interpret this to implicitly mean that features 1b and 1c are unlikely.
- They indicate that less differentiate cells have the capacity to self-renew, which implies that proliferation is symmetric and that feature 3a is unlikely.
- Naïve cells are shown to have high proliferative capacity, so this suggests support for feature 3c and implies some support for feature 3b.
- Effector memory cells (which we assume to be effector-like) are shown to have some capacities to proliferate and persist, so we assume that features 3b and 3d are implicitly (or somewhat) likely.

Xu *et al.* (2015) [24] present a pathway for differentiation:

- They propose a linear pathway from memory to effector cells, which suggests that features 1a and 1d are unlikely.
- They do not suggest branching or bidirectional differentiation pathways, so we interpret this to implicitly mean that features 1b and 1c are unlikely.
- They suggest that memory cells develop upon antigenic stimulation, and afterwards proceed to further differentiate, which implies that effector differentiation does not require antigenic stimulation and that feature 1f is likely.
- The pathway diagram indicates that memory cells self-renew (symmetrically proliferate), effector

cells have some capacity to do so, and naïve cells do not, which suggests that features 3a and 3c are unlikely, and feature 3b is implicitly (or somewhat) likely.

Golubovskaya and Wu (2016) [25] suggest separate differentiation pathways for CD4$^+$ and CD8$^+$ T cells, which we consider equally here:

- They propose a pathway for CD4$^+$ T cells that is bidirectional between effector and effector memory cells which somewhat supports feature 1c (does not fully support feature 1c, since these compartments are both considered effector-like in our framework, however, it is a related concept)
- They also propose a linear pathway from memory to effector CD8$^+$ T cells, which suggests that features 1a and 1d are unlikely.
- They do not suggest a branching differentiation pathway, so we interpret this to implicitly mean that feature 1b is unlikely.
- For each pathway, they indicate that only the first differentiation event requires stimulation by an antigen-presenting cell, which supports feature 1f.
- They indicate that naïve cells have high proliferative potential and effector cells have some potential (at least effector memory cells do), which suggests that feature 3b is implicitly (or somewhat) likely and feature 3c is likely.

Chan *et al.* (2021) [26] propose pathways for acute and chronic infection:

- They suggest that naïve cells differentiate to memory and effector cells, but the pathway diagram does not explicitly represent branching differentiation, so we assume that feature 1b is implicitly likely and that other features relating to differentiation are unclear (not represented in Table 2 of the main text).
- They depict T cell dysfunction in a pathway diagram but do not indicate back-differentiation from dysfunctional cells to memory or effector cells, which implies that feature 1g is unlikely.
- They depict that dysfunctional cells arise from pools of naïve, memory and effector cells but do not specify which cell subtype contributes to the differentiation of dysfunctional cells, so we assume that features 2a, 2b and 2c are neutral.
- The pathways indicate self-renewal (symmetric proliferation) of memory cells, do not indicate naïve proliferation, and indicate less significant proliferation of effector memory cells but not short-lived effector cells, which suggests that feature 3a is unlikely, feature 3b is neutral and feature 3c is implicitly unlikely.
- They suggest that short-lived effector cells (prone to cell death) may become long-lived effector cells (not prone to cell death), which suggests neutral support for feature 3d.

Gumber and Wang (2022) [27] suggest a pathway involving differentiation and dysfunction:

- They suggest that naïve cells differentiate to effector cells which then differentiate to memory cells after antigen clearance (not necessarily the case during short-term *ex vivo* expansion), so we assume that feature 1a is implicitly likely and feature 1d is implicitly unlikely.
- Since they report that naïve cells differentiate directly to effector cells, this indicates that they do not support branching differentiation (feature 1b).
- Since memory cells are stated to arise only after antigen is cleared, this suggests that effector cells require little antigen to differentiate to memory cells, so feature 1e is implicitly likely.
- They suggest that low levels of dysfunction may be reversible, which implies support for feature 1g.
- They report that effector cells become dysfunctional cells, and that terminal dysfunctional cells are prone to cell death, which suggests that features 2c and 3e are likely.
- They report that effector cells die after antigen clearance, which indicates that effector cells die but perhaps not during short-term *ex vivo* expansion, hence feature 3d is implicitly (or somewhat) likely.

Bulliard *et al.* (2023) [28] present pathways for differentiation and dysfunction under acute and chronic stimulation:

- The pathway indicates branching differentiation from naïve cells to memory precursor cells and short-lived effector cells (which we assume to have memory-like and effector-like phenotypes respectively) and that memory cells differentiate to effector cells, which indicates that features 1a

and 1d are unlikely and feature 1b is likely.
- They do not suggest a bidirectional differentiation pathway, so we interpret this to implicitly mean that feature 1c is unlikely.
- They suggest that T cell dysfunction cannot be reversed, hence feature 1g is unlikely.
- The pathway shows that memory precursor cells become dysfunctional, and it does not show that naïve or effector cells do so, which implicitly suggests that features 2a and 2c are unlikely, and supports feature 2b.
- They state that T cells gradually lose the ability to proliferate as they transition to effector cells, which implies that feature 3b is unlikely.
- They indicate that effector and dysfunctional cells have low persistence, which implicitly supports features 3d and 3e.

Chow *et al.* (2022) [29] discuss T cell dysfunction:
- They suggest that dysfunction may be partially reversed, which implies some support for feature 1g.
- They state that naïve cells differentiate along a range of dysfunctional cell states, which supports feature 2a.
- They suggest that overstimulation of functional cells can result in cell death, which implies support for feature 3d.
- Dysfunctional cells are said to be prone to overstimulation-induced cell death, but also that this kind of dysfunction (exhaustion) may have evolved to mitigate cell death, so we assume that feature 3e is neutral.

Zhang *et al.* (2023) [30] present experimental data regarding the *ex vivo* expansion of memory, effector and dysfunction-associated T cells:
- They indicate that the differentiation status of cells expanded *ex vivo* increases from memory-like to effector-like phenotypes, which implies that features 1a and 1d are unlikely.

Chen *et al.* (2024) [35] present a pathway for differentiation:
- They present a linear pathway from memory to effector cells, which suggests that features 1a and 1d are unlikely.
- They do not suggest branching or bidirectional differentiation pathways, so we interpret this to mean that features 1b and 1c are implicitly unlikely.
- They indicate that cells self-renew, which we assume to mean symmetric proliferation and that feature 3a is unlikely.
- Naïve cells are shown to have high proliferative potential, whereas effector-like cells have very little proliferative potential, therefore feature 3b is implicitly unlikely and feature 3c is likely.
- They indicate that effector cells have low persistence, which implies that they may be more prone to death, so feature 3d is implicitly likely.

Zeng *et al.* (2020) [56] present a pathway involving both differentiation and dysfunction under acute and chronic immune stimulation:
- Under acute stimulation, naïve cells may differentiate to memory or effector cells, and memory cells differentiate to effector cells, which supports feature 1b, and suggests that features 1a and 1d are unlikely.
- They do not suggest a bidirectional differentiation pathway, so we interpret this to implicitly mean that feature 1c is unlikely.
- They suggest that there is an unknown conversion between early dysfunctional cells and memory cells, and similarly for conversions between early dysfunctional cells and effector cells, which implies that features 1g, 2b and 2c are undecided/neutral.
- The pathway indicates that naïve cells become dysfunctional which supports feature 2a.
- They state that naïve cells proliferate in the early stages of infection, which we assume supports feature 3c.

Wherry (2011) [57] proposes a pathway for differentiation and dysfunction:
- They state that naïve cells first differentiate to effector cells and then effector cells differentiate

to memory cells after antigen clearance (which may not necessarily be the case during short-term expansion), which implies support for feature 1a and against feature 1d, and suggests that feature 1b is unlikely.
- They suggest a pathway that does not involve bidirectional differentiation between memory and effector cells, so we interpret this to mean that feature 1c is implicitly unlikely.
- Since memory cells are stated to arise only after antigen is cleared, this suggests that effector cells require little antigen to differentiate to memory cells, so feature 1e is implicitly likely.
- They suggest that early dysfunctional T cells may be able to back-differentiate to effector cells under specific circumstances, which implies support for feature 1g.
- The pathway shows that effector cells become dysfunctional but no pathways from naïve or memory cells to dysfunctional cells are shown, which suggests that features 2a and 2b are implicitly unlikely and feature 2c is likely.
- They indicate that dysfunctional cells may die, which supports feature 3e.

Baessler and Vignali (2024) [58] present a pathway for differentiation and dysfunction:
- Memory cells are shown to differentiate into effector cells, which suggests that features 1a and 1d are unlikely.
- They report that naïve T cells may differentiate into memory cells or effector cells after activation, which somewhat supports feature 1b.
- Their suggested pathway does not indicate bidirectional differentiation, but they state that effector cells differentiate back into memory cells following antigen clearance, so we assume that feature 1c is implicitly likely.
- They suggest that initial differentiation occurs after antigenic stimulation, and that memory cells may then differentiate to effector cells on antigen rechallenge, which implies that feature 1f is unlikely (both differentiation events require antigenic stimulation).
- Their pathway does not indicate that dysfunctional cells can differentiate back to a cell with a functional phenotype, and they state that terminal dysfunction cannot be easily reversed, but early dysfunctional cells can be targeted. Overall, we assume this to mean that feature 1g is implicitly unlikely (dysfunction-associated cells do not back-differentiate).
- The pathway suggests that naïve cells may also become dysfunctional after activation, which suggests that feature 2a is implicitly likely.
- No pathway from memory or effector cells to dysfunctional cells is shown, which implicitly suggests that features 2b and 2c are unlikely.
- Memory cells are shown to self-renew (symmetric proliferation), which suggests that feature 3a is unlikely.
- Effector cells are stated to die as the immune challenge resolves, which indicates that effector cells die but perhaps not during short-term *ex vivo* expansion, hence feature 3d is implicitly (or somewhat) likely.

Lugli and Gattinoni (2015) [59] propose a pathway for differentiation and briefly discuss dysfunction:
- They present a pathway depicting differentiation from memory to effector cells and transitions between activated states, which suggests that features 1a and 1d are unlikely.
- The transitions between activated cell states seems to provide the possibility for branching differentiation from naïve cells to memory or effector cells (though this is not stated), so we assume that this suggests neutral support for feature 1b.
- They do not suggest a bidirectional differentiation pathway, so we interpret this to implicitly mean that feature 1c is unlikely.
- They state that the duration of antigenic stimulation dictates the level of differentiation, which supports feature 1e (differentiation to memory cells increases when there is less stimulus) and implies that feature 1f is unlikely.
- They state that dysfunctional cells may be rejuvenated, which suggests that they may be able to regain effector function, implying support for feature 1g.
- Memory cells are stated to self-renew, which we assume indicates symmetric proliferation, and effector cells are shown to have very little to no proliferative ability, which suggests that feature 3a is implicitly unlikely and feature 3b is unlikely.
- Naïve cells are stated to have high proliferative capacity after activation, which somewhat sup-

ports feature 3c.
- Terminal effector cells are stated to be more prone to cell death than memory cells, which implies support for feature 3d.

Kallies *et al.* (2020) [61] discuss T cell dysfunction and the potential for reversing dysfunction:
- They seem to suggest that reversing dysfunction may be possible in the future but that more work is needed to determine the means to do so, which implies that feature 1g is unlikely for current experiments.
- They state that dysfunction occurs once effector cells are expanded, which implies that effector cells become dysfunctional (feature 2c is implicitly likely).

Chen *et al.* (2023) [64] present pathways for differentiation and dysfunction in acute and chronic infections:
- They present a pathway for acute infection including branching differentiation from naïve cells to memory precursor cells and short-lived effector cells (which we assume to have memory-like and effector-like phenotypes respectively) and differentiation from memory precursor to effector cells, with supports feature 1b and suggests that features 1a and 1d are unlikely.
- They do not suggest a bidirectional differentiation pathway, so we interpret this to implicitly mean that feature 1c is unlikely.
- They report that long-term antigen exposure irreversibly hinders T cell functions, but that cytokine signalling may help to reverse dysfunction, so we assume that feature 1g is neutral.
- They show that naïve cells become dysfunctional, but they do not indicate that memory or effector cells become dysfunctional, which suggests that feature 2a is likely, and implies that features 2b and 2c are unlikely.
- The pathway shows that proliferative potential is highest in the effector phase, which suggests that feature 3b is likely.

Gebhardt *et al.* (2023) [65] present multiple differentiation pathways for acute and chronic infections, including dysfunction:
- They present a branching pathway and suggest that memory cells differentiate to effector cells, which supports feature 1b and suggests that features 1a and 1d are unlikely.
- They do not suggest a bidirectional differentiation pathway, so we interpret this to implicitly mean that feature 1c is unlikely.
- They state that early dysfunctional cells only generate dysfunctional progeny and maintain their dysfunctional state even after antigen removal, which implies that feature 1g is unlikely.
- Naïve cells are shown to become dysfunctional and they state that there is evidence for memory cells giving rise to dysfunctional cells, which suggests that features 2a and 2b are likely.
- Memory cells are stated to self-renew, which we assume indicates symmetric proliferation, and effector cells are stated to have a low proliferative potential, which implies that features 3a and 3b are unlikely.
- Effector and dysfunctional cells are shown to die, which supports features 3d and 3e.

Sun *et al.* (2023) [66] propose separate pathways for T cells under acute and chronic stimulation:
- The pathway for an acute infection shows that naïve cells may differentiate to memory and effector cells, and then effector cells differentiate to memory cells during the contraction phase, which implies support for feature 1a and against feature 1d (this differentiation may not necessarily occur during short-term *ev vivo* expansion), and supports feature 1b.
- They suggest a pathway that does not involve bidirectional differentiation between memory and effector cells, so we interpret this to mean that feature 1c is implicitly unlikely.
- They state that T cell dysfunction may not be reversed, which suggests that feature 1g is unlikely.
- The pathway for chronic stimulation shows that memory precursor cells (which we assume to be memory-like) differentiate to early dysfunctional cells but they do not indicate that naïve or effector cells become dysfunctional, which supports feature 2b and implies that features 2a and 2c are unlikely.
- The pathway indicates self-renewal (symmetric proliferation) of the memory cell population but do not indicate naïve or effector cell proliferation, which we reason indicates that feature 3a is

unlikely and features 3b and 3c are implicitly unlikely.

- They indicate that effector cells die after antigen clearance (which does not necessarily occur during *ex vivo* expansion), so we assume that this means feature 3d is implicitly likely.

Larbi and Fulop (2014) [67] present a differentiation pathway and discuss dysfunction:

- A linear differentiation pathway from memory to effector cells is shown, hence features 1a and 1d are unlikely.
- They hypothesize that naïve T cells may also differentiate directly into effector cells, however, they also provide other hypotheses, so feature 1b is undecided/neutral.
- They do not suggest a bidirectional differentiation pathway, so we interpret this to implicitly mean that feature 1c is unlikely.
- They state that exhaustion is reversible, which implies support for feature 1g.
- They state that only effector-like cells express PD-1 (marker associated with T cell exhaustion) which indicates that effector cell dysfunction is more significant than naïve and memory cell dysfunction, hence features 2a and 2b are implicitly unlikely and feature 2c is implicitly likely.
- They propose that a type of naïve T cell undergoes proliferation, which supports feature 3c.

Kaech and Cui (2012) [68] present multiple potential pathways for differentiation:

- They do not suggest that one pathway is more valid than another, so we interpret this to mean that they have neutral opinions for features 1a, 1b, 1d and 1e (linear differentiation from memory to effector cells, and a branching model with no differentiation between memory and effector cells and where memory cell differentiation arises with less stimulus).
- They do not suggest a bidirectional differentiation pathway, so we interpret this to implicitly mean that feature 1c is unlikely.
- In each pathway, antigenic stimulation is required for all differentiation events, which suggests that feature 1f is unlikely.
- Some differentiation pathways indicate self-renewal (symmetric proliferation) of memory cells, and one indicates the possibility for asymmetric division, so we assume that feature 3a is undecided/neutral.
- Effector cells are not shown to proliferate, so we assume that feature 3b is implicitly unlikely.
- Naïve cells are not shown to proliferate in most pathways, but are shown to proliferate in the asymmetric division model, which suggests that feature 3c is undecided/neutral.
- Effector cells are shown to die in all pathways, which suggests that feature 3d is likely.

June (2007) [69] presents multiple potential differentiation pathways:

- They do not suggest that one pathway is more valid than another, so we interpret this to mean that they have neutral opinions for features 1a, 1b, 1e and 1f (linear differentiation from naïve to effector to memory cells where only naïve to effector differentiation requires antigen, and a branching model in which memory differentiation is supported by less stimulus and central memory cells differentiate to effector memory cells).
- They do not suggest a bidirectional differentiation pathway, so we interpret this to implicitly mean that feature 1c is unlikely.
- All pathways suggest some differentiation between memory and effector cells, which suggests that feature 1d is unlikely.
- They indicate that effector cells can die, which supports feature 3d.

Raphael *et al.* (2020) [70] present multiple potential differentiation pathways:

- They do not suggest that one pathway is more valid than another, so we interpret this to mean that they have neutral opinions for features 1a, 1b, 1d and 3a (models with differentiation from effector cells to memory cells, asymmetric division/branching without differentiation between memory and effector cells, and some that indicate self renewal/symmetric proliferation).
- They do not suggest a bidirectional differentiation pathway, so we interpret this to implicitly mean that feature 1c is unlikely.
- They state that memory cell differentiation inversely correlates with stimulation strength, which suggests that feature 1e is likely.
- Each potential pathway indicates that only naïve cells require antigenic stimulation, which implies

support for feature 1f.
- Some pathways indicate that memory cells proliferate but not effector cells which implies feature 3b is unlikely.

Yi *et al.* (2018) [71] present a pathway for T cell differentiation:
- They present a branching pathway with no differentiation between memory and effector cells, which supports features 1b and 1d.
- They do not suggest a bidirectional differentiation pathway, so we interpret this to implicitly mean that feature 1c is unlikely.
- The pathway depicts self-renewal (symmetric proliferation) of memory cells, but does not indicate that naïve or effector cells proliferate, which suggests that feature 3a is unlikely and implies that features 3b and 3c are unlikely.
- The pathway depicts the death of effector memory cells (which we assume to have effector-like phenotypes), which supports feature 3d.

Espinosa *et al.* (2016) [72] present multiple potential pathways for differentiation:
- Each pathway depicts differentiation from effector cells to memory cells, which suggests that feature 1a is likely and feature 1d is unlikely.
- They do not suggest that one pathway is more valid than another, so we interpret this to mean that they have neutral opinions for features 1b and 1c (branching and bidirectional differentiation).

Sun *et al.* (2024) [73] present a pathway involving differentiation and dysfunction:
- They present a branching pathway and suggest that memory cells progressively differentiate to effector cells, which supports feature 1b and suggests that features 1a and 1d are unlikely.
- They do not suggest a bidirectional differentiation pathway, so we interpret this to implicitly mean that feature 1c is unlikely.
- They present a pathway involving dysfunction but do not indicate differentiation from dysfunctional cells back to effector cells, so we assume that this means feature 1g is implicitly unlikely.
- The pathway shows naïve cells differentiating into early dysfunctional cells and effector memory cells (which we assume to have effector-like phenotypes) differentiating into terminal dysfunctional cells, which supports features 2a and 2c.
- Stem memory cells are shown to differentiate into early dysfunctional cells with a dashed arrow, which indicates that feature 2b is implicitly (or somewhat) likely.
- The pathway depicts self-renewal (symmetric proliferation) of memory and early dysfunctional cells, but not naïve or effector cells, which suggests that feature 3a is unlikely and implies that features 3b and 3c are unlikely.

Kartikasari *et al.* (2019) [76] present a pathway for $CD8^+$ T cell differentiation and dysfunction:
- The pathway includes branching differentiation from naïve cells to memory and effector cells with further differentiation from effector cells to memory cells, which indicates that features 1a and 1b are likely, and feature 1d is unlikely.
- They suggest a pathway that does not involve bidirectional differentiation, so we interpret this to mean that feature 1c is implicitly unlikely.
- They report that dysfunction can be reversed, which implies support for feature 1g.
- The pathway shows that effector cells become dysfunctional but no pathways from naïve or memory cells to dysfunctional cells are shown, which implies that features 2a and 2b are unlikely and suggests that feature 2c is likely.

Kaech *et al.* (2002) [77] present multiple potential differentiation pathways, with one involving dysfunction:
- They do not suggest that one pathway is more valid than another, so we interpret this to mean that they have neutral opinions for features 1b and 1d (linear differentiation or branching without differentiation between memory and effector cells).
- They do not suggest a bidirectional differentiation pathway, so we interpret this to implicitly mean that feature 1c is unlikely.

- Each pathway indicates that only the first differentiation event requires stimulus and most indicate that effector cells differentiate to memory cells, which indicates support for features 1a and 1f.
- They propose a pathway in which effector cells become dysfunctional and then die, which indicates that features 2c and 3e are likely, and they do not indicate any back-differentiation from dysfunctional cells to effector cells, which suggests that feature 1g is implicitly unlikely.
- They state that memory cells self renew, which we assume to indicate symmetric proliferation and that feature 3a is implicitly unlikely.
- They state that effector cells have a limited proliferative potential compared with naïve and memory cells, which implies that naïve cells proliferate but that effector cell proliferation is less significant, so we assume feature 3b is implicitly unlikely and feature 3c is implicitly likely.
- They suggest that effector cells die during the contraction phase, which indicates that effector cells die but perhaps not during short-term *ex vivo* expansion, hence feature 3d is implicitly (or somewhat) likely.

Borsa *et al.* (2019) [78] discuss asymmetric division in the context of memory T cells and antigenic stimulation:

- They state that naïve cells first differentiate to effector cells and then effector cells differentiate to memory cells after antigen clearance, which implies supports for feature 1a and against feature 1d (this differentiation may not necessarily occur during short-term *ev vivo* expansion) and does not support feature 1b.
- Since memory cells are stated to arise only after antigen is cleared, this suggests that effector cells require little antigen to differentiate to memory cells, so feature 1e is implicitly likely.
- They report that effector functions may be restored in dysfunctional cells, which supports feature 1g.
- Especially in environments with strong stimulus, naïve and memory cells can undergo asymmetric division, and they also state that effector cells can proliferate symmetrically, which supports features 3a, 3b and 3c.
- They report that effector cells die after antigen clearance, which indicates that effector cells die but perhaps not during short-term *ex vivo* expansion, hence feature 3d is implicitly (or somewhat) likely.

Wherry *et al.* (2004) [79] propose a pathway for differentiation and dysfunction in acute and chronic infection:

- They state that naïve cells differentiate into effector cells and then into memory cells after antigen clearance, which implies that feature 1a is likely and feature 1d is unlikely (this differentiation may not necessarily occur during short-term *ev vivo* expansion) and suggests that feature 1b is unlikely.
- They do not suggest a bidirectional differentiation pathway, so we interpret this to implicitly mean that feature 1c is unlikely.
- They report that rest from antigenic stimulation is critical for memory cell differentiation, which implies that feature 1e is likely (differentiation to memory cells increases when there is less stimulus).
- They report that chronic memory (or early dysfunctional) cells may not be reverted to a more functional state, which implies that feature 1g is unlikely.
- The pathway shows effector cells becoming dysfunctional, and they do not show a pathway from naïve or normal memory cells to dysfunctional cells, which suggests that features 2a and 2b are implicitly unlikely, and feature 2c is likely.
- The pathway indicates self-renewal (symmetric proliferation) of memory cells and they state that naïve cells proliferate, but they do not indicate effector cell proliferation, which suggests that feature 3a is unlikely, feature 3b is implicitly unlikely, and feature 3c is likely.
- They report that chronic memory (or early dysfunctional) cells have limited persistence (susceptible to cell death) after antigen clearance (not necessarily the case during short-term expansion), so we assume that feature 3e is implicitly likely.

Kirman *et al.* (2017) [80] present multiple potential differentiation pathways and comment on dysfunction:

- They do not suggest that one pathway is more valid than another, so we interpret this to mean that they have neutral opinions for features 1a, 1b, 1d and 1e (linear differentiation from effector cells to memory cells, a branching model in which memory cell differentiation is supported with a lower level of activation and with memory cells differentiating to effector cells, and a branching model without differentiation between memory and effector cells).
- They suggest that the pathways they proposed may co-exist, which allows for the possibility of bidirectional differentiation, hence feature 1c is also neutral.
- They state that prolonged antigen exposure can lead to exhausted effector cells, which implies support for feature 2c.
- Each pathway indicates self-renewal (symmetric proliferation) of the memory cell population but one depicts asymmetric division, so we assume that feature 3a is undecided/neutral.
- They do not indicate effector cell proliferation, which we reason indicates that feature 3b is implicitly unlikely.
- The asymmetric division model suggests that naïve cells proliferate but the other models do not, which indicates neutral support for feature 3c.
- All pathways explicitly indicate effector cell death, and they indicate that cell death may occur alongside dysfunction, which supports feature 3d and implies support for feature 3e.

Mahnke *et al.* (2013) [81] propose a pathway for differentiation and comment on dysfunction:

- They suggest a pathway in which cells transition between activated states, generally trending from memory cells to effector cells (without direct differentiation between these discrete subtypes), which implies that features 1a and 1d are unlikely.
- The transitions between activated cell states seems to provide the possibility for branching differentiation from naïve cells to memory or effector cells, so we assume that this suggests neutral support for feature 1b.
- They do not suggest a bidirectional differentiation pathway, so we interpret this to implicitly mean that feature 1c is unlikely.
- They indicate that memory cells are less dependent on antigen and effector cells are more dependent, which implies that feature 1e is likely.
- They state that cells become dysfunctional at the effector stage, memory cells express markers associated with exhaustion (though not as much as effector cells), but naïve cells are not indicated to express these markers, which suggests that feature 2a is implicitly unlikely, feature 2b is implicitly likely, and feature 2c is likely.
- Memory cells are stated to self-renew, which we assume indicates symmetric proliferation, and naïve and effector cells are shown to have little proliferative potential (and do not typically express markers for proliferation), which indicates that feature 3a is implicitly unlikely and features 3b and 3c are unlikely.

Fazeli *et al.* (2023) [82] propose multiple pathways for T cell differentiation and comment on dysfunction:

- They propose a linear pathway from memory to effector cells in which memory cell differentiation is supported by less stimulation, but they state that this model is in conflict with the definition of stem memory cells, which implies that features 1a and 1e are unlikely.
- They also propose a branching model without differentiation between memory and effector cells, without suggesting that one is more likely than the other, so we assume that they have neutral opinions for features 1b, 1c, and 1d.
- Each proposed pathway suggests that only the first differentiation event requires antigenic stimulation, which supports feature 1f.
- They state that stem memory cells are the least exhausted subtype compared to other memory subpopulations, which implies that feature 2b is unlikely.
- They discuss the possibility of naïve T cells proliferating asymmetrically without certainty, which somewhat supports (or implies support for) features 3a and 3c.
- They present a diagram which depicts effector cells proliferating, hence feature 3b is likely.

Liisborg *et al.* (2021) [83] present a differentiation pathway:

- They propose a linear differentiation pathway from memory to effector cells, which suggests that features 1a and 1d are unlikely.
- They do not suggest branching or bidirectional differentiation pathways, so we interpret this to implicitly mean that features 1b and 1c are unlikely.
- Naïve cells and effector memory cells (assumed to be effector-like) are shown to have high proliferative potential, but terminal effector memory cells do not proliferate, which implies support for feature 3b and supports feature 3c.

Lan *et al.* (2023) [84] present pathways for differentiation and dysfunction during acute and chronic stimulation:
- They present a branching pathway and suggest that memory cells differentiate to effector cells, which supports feature 1b and suggests that features 1a and 1d are unlikely.
- They do not suggest a bidirectional differentiation pathway, so we interpret this to implicitly mean that feature 1c is unlikely.
- The pathway indicates that antigen is required for each stage of differentiation from naïve to memory to effector cells, which suggests that feature 1f is unlikely.
- The pathway shows that early dysfunctional cells differentiate to effector-like cells, which supports feature 1g.
- In the pathway, naïve cells are shown to become early dysfunctional cells, memory cells are indicated to have an unknown transition to dysfunctional cells, and effector cells are not shown to become dysfunctional, which we assume supports feature 2a, suggests undecided/neutral support feature 2b, and implies that feature 2c is unlikely.
- The pathway indicates self-renewal (symmetric proliferation) of memory and early dysfunctional cells but not naïve or effector cells, which suggests that feature 3a is unlikely and features 3b and 3c are implicitly unlikely.
- They report that dysfunctional cells do not simply die, which implies that feature 3e is unlikely.

Gräbnitz *et al.* (2023) [85] discuss asymmetric proliferation in the context of antigenic stimulation:
- They suggest that naïve cells proliferate asymmetrically (especially with large stimulus presence such as during *ex vivo* expansion) into early memory and effector cell subtypes, which also suggests a branching pathway, so we assume that feature 1b is implicitly likely and features 3a and 3c are likely.
- They also suggest that strong stimulation favours effector cell differentiation and weak stimulation favours memory cell differentiation, which supports feature 1e.

King *et al.* (2012) [86] discuss differentiation and the possibility of asymmetric cell division:
- They suggest that effector cell differentiation is favoured by strong stimulation, which implies support for feature 1e.
- They support the idea of asymmetric naïve cell proliferation (especially with large stimulus presence such as during *ex vivo* expansion), which also suggests a branching pathway, so we assume that feature 1b is implicitly likely and features 3a and 3c are likely.

Jia *et al.* (2024) [87] discuss T cell dysfunction and the potential to regain effector function in early dysfunctional cells:
- They suggest that early dysfunctional cells may back-differentiate to effector cells, which supports feature 1g.
- They also suggest that naïve cells can differentiate to early dysfunctional cells, which supports feature 2a.
