## Supplementary Text 2 for "A computational investigation of pathways for *ex vivo* differentiation and dysfunction of healthy and patient T cells"

ORCIDs: MSL, 0009-0009-6633-2044; YB, 0000-0001-9901-7782; PRB, 0000-0003-3962-5393; ALJ, 0000-0001-9103-7092

October 19, 2026

Table S2.1: **Marker expressions to model T cell subtypes.** Expressions that we use to identify T cell phenotypic compartments, where + indicates that the modelled subtype expresses the marker, and − indicates that it does not. In the PD-1/TIM-3 column, the first marker indicates PD-1 expression and the second indicates TIM-3 expression. Blank cells indicate that the marker was not considered for that cell type (may be + or −).

| T cell subtype | CD45RA | CCR7 | CD95 | CD25 | PD-1/TIM-3 | Ref. |
|---|---|---|---|---|---|---|
| Naïve | + | + | − | − | −/−, +/−, or −/+ | [2, 3] |
| Memory stem (memory-like) | + | + | + | + | −/−, +/−, or −/+ | [2] |
| Central memory (memory-like) | − | + | | | −/−, +/−, or −/+ | [2] |
| Effector memory (effector-like) | − | − | | | −/−, +/−, or −/+ | [2] |
| Terminal effector (effector-like) | + | − | | | −/−, +/−, or −/+ | [2] |
| Dysfunction-associated | | | | | +/+ | [4] |

### S2 Text. Calculation of cell counts from data

Data published by Zhang *et al.* [1] describes the proportions of expanded T cells that express various markers relevant to differentiation and dysfunction, namely CD45RA, CCR7, CD25, CD95, PD-1 and TIM-3. They provide these proportions for T cells that express CD4 and for those that express CD8, and CD95 expression data is only provided for the initial population of cells. Table S2.1 summarises the expressions that we used to identify each relevant T cell compartment for the purpose ofn modelling (Table 1 of the main text), but we note that cells in each compartment may also express markers other than those indicated in this table. For example, naïve cells are reported to co-express CD45RA and CCR7 but they do not express CD95, whereas memory stem cells are reported to express CD45RA, CCR7 and CD95 [2]. While some naïve cells may express CD25, it is commonly thought to be an indicator for activation and the majority of naïve cells typically do not express CD25 [3]. Although CD25 expression is not typically used as an indicator for distinguishing naïve and memory stem cells, we do so due to a lack of data on CD95 expression at Day 8.

Following Zhang *et al.* [1], we use PD-1 and TIM-3 co-expression as an operational marker of an exhaustion-associated phenotype [4]. Although PD-1/TIM-3 co-expression alone does not establish terminal or irreversible dysfunction, in our reduced model, this phenotype is assigned to the dysfunction-associated compartment. To ensure that each cell subtype is functionally distinct, we assume that this dysfunctional state is exclusive such that naïve, memory-like and effector-like cells may not co-express PD-1 and TIM-3. We acknowledge that other markers could have been used to identify the relevant T cell phenotypes, so our selection is a modelling assumption.

Let the day $n$ proportion of CD4$^+$ cells expressing marker X and not expressing marker Y be $P^{(n)}_{\text{X}^+\text{Y}^-}$ (for example, the day 0 proportion of cells expressing CD25 is $P^{(0)}_{\text{CD25}^+}$, and the Day 8 proportion of cells expressing CD45RA but not CCR7 is $P^{(8)}_{\text{CD45RA}^+\text{CCR7}^-}$), which is estimated from the data. Following Table S2.1, for each experiment (8 samples of healthy donor T cells, 10 samples of patient T cells, each expanded under 7 levels of stimulation), we compute the proportion of CD4$^+$ cells that are naïve $P_\text{N}$, memory-like $P_\text{M}$ (memory stem and central memory cells), effector-like $P_\text{E}$ (effector memory and terminal effector cells) and dysfunction-associated $P_\text{D}$ cells at day 0 as

$$\begin{aligned}
P^{(0)}_\text{N} &= P^{(0)}_{\text{CD45RA}^+\text{CCR7}^+\text{CD95}^-}\left(1 - P^{(0)}_{\text{PD-1}^+\text{TIM-3}^+}\right),\\
P^{(0)}_\text{M} &= \left(P^{(0)}_{\text{CD45RA}^+\text{CCR7}^+\text{CD95}^+} + P^{(0)}_{\text{CD45RA}^-\text{CCR7}^+}\right)\left(1 - P^{(0)}_{\text{PD-1}^+\text{TIM-3}^+}\right),\\
P^{(0)}_\text{E} &= \left(P^{(0)}_{\text{CD45RA}^-\text{CCR7}^-} + P^{(0)}_{\text{CD45RA}^+\text{CCR7}^-}\right)\left(1 - P^{(0)}_{\text{PD-1}^+\text{TIM-3}^+}\right),\\
P^{(0)}_\text{D} &= P^{(0)}_{\text{PD-1}^+\text{TIM-3}^+},
\end{aligned}$$

and at day 8 as

$$
\begin{aligned}
P_{\mathrm{N}}^{(8)} &= P_{\mathrm{CD45RA^+CCR7^+}}^{(8)} P_{\mathrm{CD25^-}}^{(8)} \left(1 - P_{\mathrm{PD\text{-}1^+TIM\text{-}3^+}}^{(8)}\right), \\
P_{\mathrm{M}}^{(8)} &= \left(P_{\mathrm{CD45RA^+CCR7^+}}^{(8)} P_{\mathrm{CD25^+}}^{(8)} + P_{\mathrm{CD45RA^-CCR7^+}}^{(8)}\right) \left(1 - P_{\mathrm{PD\text{-}1^+TIM\text{-}3^+}}^{(8)}\right), \\
P_{\mathrm{E}}^{(8)} &= \left(P_{\mathrm{CD45RA^-CCR7^-}}^{(8)} + P_{\mathrm{CD45RA^+CCR7^-}}^{(8)}\right) \left(1 - P_{\mathrm{PD\text{-}1^+TIM\text{-}3^+}}^{(8)}\right), \\
P_{\mathrm{D}}^{(8)} &= P_{\mathrm{PD\text{-}1^+TIM\text{-}3^+}}^{(8)}.
\end{aligned}
$$

Note that we use data regarding the co-expression of CD45RA/CCR7/CD95 to compute the naïve and memory stem cell counts at day 0, but data regarding the co-expression of CD45RA/CCR7/CD25 was not available for Day 8 (only separate measurements for co-expression of CD45RA/CCR7 and expression of CD25). Therefore, we assume that $P_{\mathrm{CD45RA^+CCR7^+CD25^-}}^{(8)} \approx P_{\mathrm{CD45RA^+CCR7^+}}^{(8)} P_{\mathrm{CD25^-}}^{(8)}$, and similarly for co-expressions of PD-1/TIM-3 with other markers. By construction, the estimated proportions satisfy $P_{\mathrm{N}} + P_{\mathrm{M}} + P_{\mathrm{E}} + P_{\mathrm{D}} = 1$ at both time points.

The total number of $CD4^+$ T cells initially is $T_{\mathrm{init}} = 10^5 P_{\mathrm{CD4^+}}^{(0)}$ [1], and let the fold expansion of $CD4^+$ T cells at day 8 be $T_{\mathrm{exp}}$ for a given experiment (determined from the data). The total number of $CD4^+$ naïve cells at day 0 is then $T_{\mathrm{init}} P_{\mathrm{N}}^{(0)}$ and the total number at day 8 is $T_{\mathrm{init}} T_{\mathrm{exp}} P_{\mathrm{N}}^{(8)}$, and similarly for the other cell compartments. This process is repeated to compute the subpopulations of each cell compartment at days 0 and 8 for $CD8^+$ cells, and the total cells at days 0 and 8 are then the sum of those for $CD4^+$ and $CD8^+$ T cells. The quantities used to calibrate and validate our models are the averages and standard deviations of the total cell counts ($CD4^+$ and $CD8^+$) for each cell subpopulation at day 8, which are computed separately for each stimulation level over the 8 healthy donor T cell samples and the 10 patient-derived T cell samples.
