## Supplementary Text 3 for "A computational investigation of pathways for *ex vivo* differentiation and dysfunction of healthy and patient T cells"

ORCIDs: MSL, 0009-0009-6633-2044; YB, 0000-0001-9901-7782; PRB, 0000-0003-3962-5393; ALJ, 0000-0001-9103-7092

October 19, 2026

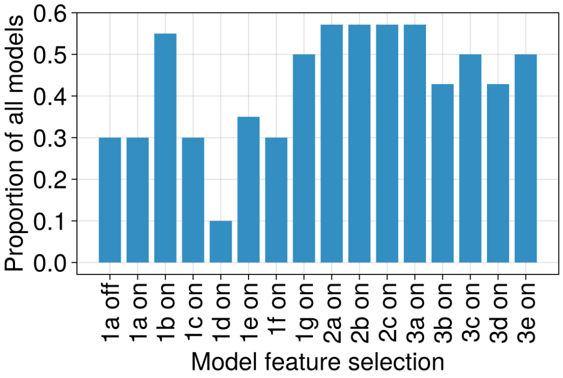


Figure S3.1: **Feature selection frequency among all simulated models.** The proportions of all simulated models (determined by the exclusions described in Table S3.1) that have each feature selection.

### S3 Text. Excluded pathway feature combinations

Although most combinations of model features are considered, some are excluded to reduce computational load due to redundancy in selected features or irrelevance of the pathways that result from specific combinations. Table S3.1 summarises the conditions and reasoning for a pathway to be excluded. Without these exclusions, $2^{15} = 32{,}768$ models may be constructed since we consider that pathways may be defined from 15 features that may be turned on or off. With these exclusions, we simulate 7,840 models, which significantly reduces computational load without losing much information about viable differentiation pathways. The frequencies of feature selections among all simulated models (none of which contain excluded feature combinations) are shown in Fig S3.1.

Despite the exclusions described in Table S3.1, some included models still exhibit combinations of features in which one feature overwrites another. Specifically, feature 1c overwrites feature 1a (naïve cells always differentiate into effector-like cells in the bidirectional model regardless of whether feature 1a is on or off), and feature 1d also overwrites feature 1a (there is no differentiation between memory-like and effector-like compartments regardless of whether feature 1a is on or off). In the cases when a model has feature 1c or 1d on, we do not consider models to have feature 1a turned off (these models are already excluded if feature 1a is on). These models may be considered to have neither feature 1a turned on nor off, hence the need to display results for feature 1a on and off (for example in Fig S3.1).

Table S3.1: **Excluded feature combinations.** Descriptions and reasoning for conditions that represent excluded combinations of feature selections (note that features turned off are represented by bars, e.g. $\overline{1a}$ represents feature 1a off).

| Condition | Description | Reason for exclusion |
|---|---|---|
| 1a and 1c | Differentiation proceeds from naïve to effector-like to memory-like states, and it is bidirectional (starting with naïve to effector). | This combination of features is redundant since differentiation already proceeds from naïve to effector-like to memory-like states in bidirectional models. When feature 1a is off and feature 1c is on, we assume the pathway to have a bidirectional structure as this also accounts for the structure normally introduced by feature 1a. |
| 1d and $\overline{1b}$ | No differentiation directly between memory-like and effector-like compartments, and differentiation is not branching. | Pathways in the literature seem to only suggests that memory-like cells do not differentiate to effector-like states when there is branching differentiation. Additionally, allowing feature 1d to be turned on without feature 1b allows for the possibility that one T cell subtype is disconnected from the pathway, which we assume to be invalid. |
| 1d and (1a or 1c or 1f) | No differentiation between memory-like and effector-like compartments, but differentiation is also linear from effector-like to memory-like states or it is bidirectional, or we consider removing stimulus dependency for differentiation between memory-like and effector-like cells. | We consider it contradictory to assert that there is no differentiation between memory-like and effector-like cells while also defining ways in which they differentiate to each other (linearly, in a bidirectional manner, or without stimulus dependency). In cases where features 1a, 1c or 1f are off, we assume that feature 1d defines the pathway structure between memory-like and effector-like cells (we remove differentiation between memory-like and effector-like compartments). |
| 1e and 1f | Rate of differentiation to memory-like states is greater when there is less stimulus, but differentiation is not dependent on stimulation after the first differentiation event. | Feature 1e asserts that differentiation to memory-like and effector-like cell states are both affected by stimulus, so removing stimulus dependency in this case does not make sense. |
| $\overline{2a}$ and $\overline{2b}$ and $\overline{2c}$ | None of naïve, memory-like or effector-like cells can upregulate inhibitory receptors and differentiate to dysfunction-associated cells. | This combination would disconnect dysfunction-associated cells from the rest of the pathway, which we assume to be invalid. |
| $\overline{3a}$ and 3b and 3d | Effector-like cells may die and divide to create two effector-like daughter cells. | This combination produces a viable differentiation pathway, but the resulting model is not structurally identifiable and fitting it to data would result in very similar population predictions as the models with features 3b or 3d turned off (just different rate of proliferation). Removing this combination reduces computational load without losing much information regarding pathways that can explain experimental data. |
