## Supplementary Text 4 for "A computational investigation of pathways for *ex vivo* differentiation and dysfunction of healthy and patient T cells"

ORCIDs: MSL, 0009-0009-6633-2044; YB, 0000-0001-9901-7782; PRB, 0000-0003-3962-5393; ALJ, 0000-0001-9103-7092

October 19, 2026

### S4 Text. Mathematical model formulation

Here we provide a detailed description of the formation of Eqs (1)–(4), which simulate the population evolution of naïve $N$, memory-like $M$, effector-like $E$, and dysfunction-associated $D$ cells (Eqs (1)–(4) of the main text). The full model is given by

$$\frac{\mathrm{d}N}{\mathrm{d}t} = \underbrace{-r_{\mathrm{m}}(s)N}_{(\overline{1\mathrm{a}}\text{ and }\overline{1\mathrm{c}})\text{ or }1\mathrm{b}} \; \underbrace{-r_{\mathrm{e}}(s)N}_{1\mathrm{a}\text{ or }1\mathrm{b}\text{ or }1\mathrm{c}} \; \underbrace{-r_{\mathrm{d}}(s)N}_{2\mathrm{a}} \; \underbrace{+\mu N}_{3\mathrm{c}} \; \underbrace{+\mu M}_{3\mathrm{a}} \; \underbrace{+\mu E}_{3\mathrm{a}\text{ and }3\mathrm{b}}\,, \tag{1}$$

$$\frac{\mathrm{d}M}{\mathrm{d}t} = \underbrace{r_{\mathrm{m}}(s)N}_{(\overline{1\mathrm{a}}\text{ and }\overline{1\mathrm{c}})\text{ or }1\mathrm{b}} \; \underbrace{-r_{\mathrm{e}}(s)M}_{(\overline{1\mathrm{a}}\text{ or }1\mathrm{c})\text{ and }\overline{1\mathrm{d}}} \; \underbrace{-r_{\mathrm{d}}(s)M}_{2\mathrm{b}} \; \underbrace{+r_{\mathrm{m}}(s)E}_{(1\mathrm{a}\text{ or }1\mathrm{c})\text{ and }\overline{1\mathrm{d}}} \; \underbrace{+\mu M}_{\overline{3\mathrm{a}}}, \tag{2}$$

$$\frac{\mathrm{d}E}{\mathrm{d}t} = \underbrace{r_{\mathrm{e}}(s)N}_{1\mathrm{a}\text{ or }1\mathrm{b}\text{ or }1\mathrm{c}} \; \underbrace{+r_{\mathrm{e}}(s)M}_{(\overline{1\mathrm{a}}\text{ or }1\mathrm{c})\text{ and }\overline{1\mathrm{d}}} \; \underbrace{-r_{\mathrm{m}}(s)E}_{(1\mathrm{a}\text{ or }1\mathrm{c})\text{ and }\overline{1\mathrm{d}}} \; \underbrace{-r_{\mathrm{d}}(s)E}_{2\mathrm{c}} \; \underbrace{+r_{\mathrm{e}}(s)D}_{1\mathrm{g}} \; \underbrace{+\mu E}_{\overline{3\mathrm{a}}\text{ and }3\mathrm{b}} \; \underbrace{-\lambda E}_{3\mathrm{d}}, \tag{3}$$

$$\frac{\mathrm{d}D}{\mathrm{d}t} = \underbrace{r_{\mathrm{d}}(s)N}_{2\mathrm{a}} \; \underbrace{+r_{\mathrm{d}}(s)M}_{2\mathrm{b}} \; \underbrace{+r_{\mathrm{d}}(s)E}_{2\mathrm{c}} \; \underbrace{-r_{\mathrm{e}}(s)D}_{1\mathrm{g}} \; \underbrace{-\lambda D}_{3\mathrm{e}}, \tag{4}$$

where $\mu$ and $\lambda$ are the constant rates of proliferation and death, respectively, and $r_{\mathrm{m}}(s)$ is the rate of differentiation to a memory-like state, $r_{\mathrm{e}}(s)$ is the rate of differentiation to an effector-like state, and $r_{\mathrm{d}}(s)$ is the rate of inhibitory receptor upregulation, which depend on the density of activating stimulus in the dish $s$.

Each ordinary differential equation in Eqs (1)–(4) consists of terms that describe one of the in-going (positive) or out-going (negative) rates for the corresponding cell compartment, as illustrated in Fig S4.1. For example, the first equation representing the naïve cell subpopulation begins with a term that describes a reduction in naïve cells due to their differentiation into memory-like cells, which is proportional to the current number of naïve cells and the rate of differentiation to memory-like phenotypes (i.e. $-r_{\mathrm{m}}(s)N$). A corresponding term begins the second equation describing the memory-like cell compartment, which describes the increase in memory-like cells due to this differentiation from naïve cells. These terms are only present in either equation if the condition written underneath them is met, namely that features 1a and 1c are both off (represented by bars above 1a and 1c, i.e. $\overline{1\mathrm{a}}$ and $\overline{1\mathrm{c}}$) or feature 1b is on.

All term conditions are summarised and justified in Table S4.1. For example, the aforementioned condition is chosen since, in our modelling framework, naïve cells will only differentiate to memory-like states if the pathway does not exhibit bidirectional differentiation (feature 1c, which we define to begin with differentiation from naïve to effector-like states) nor linear differentiation from naïve cells to effector-like cells (feature 1a), unless the pathway involves branching differentiation from naïve cells to both memory-like and effector-like cells (feature 1b).

$r_{\mathrm{m}}$ and $r_{\mathrm{e}}$ are both constant (specifically half their maximal rate) when feature 1f is on and when that differentiation does not occur directly from the naïve cell state or if there is branching differentiation (these traits were noticed in some of the literature [1–3]). That is the case for memory-like cell differentiation when features 1a, 1b or 1c are on, and for effector-like cell differentiation when features 1a and 1c are both off or feature 1b is on. Also, $r_{\mathrm{m}}$ changes to decrease as stimulus density increases when feature 1e is on (also indicated by some biological literature [4, 5]). Since the values of $r_{\mathrm{m}}$ and $r_{\mathrm{e}}$ are conditional on pathway features, these conditions must also be evaluated for a given model. In full, the rates of differentiation and dysfunction (extensions of Eqs (5)–(7) of the main text)

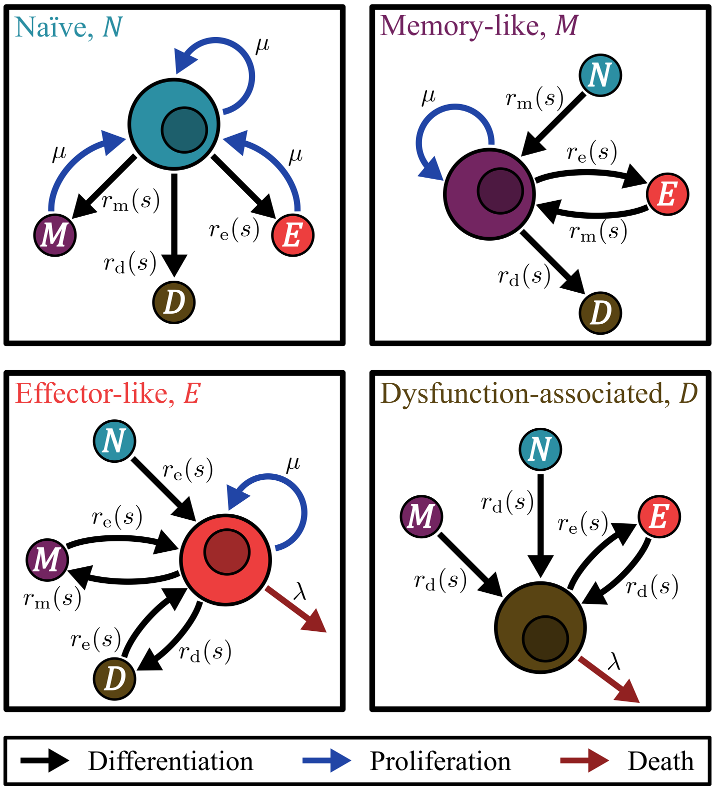


Figure S4.1: **In-going and out-going rates for each cell type.** Diagrams for naïve (blue), memory-like (purple), effector-like (red) and dysfunction-associated (brown) T cell compartments detailing the rates for transitions, proliferation and death which inform terms in Eqs (1)–(4).

are

$$r_{\mathrm{m}}(s) = \begin{cases} \dfrac{r_{\mathrm{m}}^{\max} s}{s_{\mathrm{m}} + s} & \text{if } \overline{\text{1e}} \text{ and not (1f and (1a or 1b or 1c))} \\ r_{\mathrm{m}}^{\max}\left(1 - \dfrac{s}{s_{\mathrm{m}} + s}\right) & \text{if 1e and not (1f and (1a or 1b or 1c))} \\ \dfrac{1}{2} r_{\mathrm{m}}^{\max} & \text{if 1f and (1a or 1b or 1c)} \end{cases}, \tag{5}$$

$$r_{\mathrm{e}}(s) = \begin{cases} \dfrac{r_{\mathrm{e}}^{\max} s}{s_{\mathrm{e}} + s} & \text{if not (1f and ((}\overline{\text{1a}} \text{ and } \overline{\text{1c}}\text{) or 1b))} \\ \dfrac{1}{2} r_{\mathrm{e}}^{\max} & \text{if 1f and ((}\overline{\text{1a}} \text{ and } \overline{\text{1c}}\text{) or 1b)} \end{cases}, \tag{6}$$

$$r_{\mathrm{d}}(s) = \frac{r_{\mathrm{d}}^{\max} s}{s_{\mathrm{d}} + s}, \tag{7}$$

where $r_{\mathrm{m}}^{\max}$, $r_{\mathrm{e}}^{\max}$ and $r_{\mathrm{d}}^{\max}$ are the maximum rates of differentiation to memory-like, effector-like and dysfunction-associated states (in day$^{-1}$), respectively, and $s_{\mathrm{m}}$, $s_{\mathrm{e}}$ and $s_{\mathrm{d}}$ are the half-maximum stimulus densities, which are the values of $s$ at which the differentiation rates are half their maximal values (ng of activating stimulus per µg of micro-rods). Fig S4.2 shows the rate of differentiation to memory-like cells $r_{\mathrm{m}}(s)$ for different stimulus densities under each case, using parameter values close to the median for the best patient T cell models. The base case for $r_{\mathrm{e}}$ (when feature 1f is off) and $r_{\mathrm{d}}$ take a similar form to the base case of $r_{\mathrm{m}}$, and the case with feature 1f on for $r_{\mathrm{e}}$ takes a similar form to that case of $r_{\mathrm{m}}$.

In the case studies, we presented mathematical models to approximate differentiation pathways suggested by Wherry [6] (case study 1) and Sun *et al.* [7] (case study 2). To derive these models, we reduced Eqs (1)–(4) to contain only the relevant terms by considering the features involved in these pathways and the conditions on those features in Table S4.1. Table S4.2 details the features that we selected for the pathways, which are mostly informed by the suggestions made in these articles (we

Table S4.1: **Feature conditions in the mathematical model.** Descriptions of the corresponding terms and justifications for conditions in Eqs (1)–(4) (note that features turned off are represented by bars). Features 1e and 1f are not represented in these conditions as they do not alter the structure of the pathway, rather the rates of differentiation.

| Condition | Description of term | Justification |
|---|---|---|
| ($\overline{1a}$ and $\overline{1c}$) or 1b | Differentiation from naïve to memory | This may only occur if the pathway does not exhibit linear differentiation from naïve to an effector-like state nor bidirectional differentiation starting with naïve to effector, unless there is branching differentiation |
| 1a or 1b or 1c | Differentiation from naïve to effector | This may only occur if the pathway exhibits linear differentiation from naïve to an effector-like state, bidirectional differentiation, or branching differentiation |
| 2a | Differentiation from naïve to dysfunction-associated | This may only occur if naïve cells can upregulate inhibitory receptors |
| 3c | Naïve cell proliferation | This may only occur if naïve cells can proliferate |
| 3a | Memory-like cell proliferation creating a naïve daughter cell | This may only occur when proliferating cells form new less differentiated cells, which we assume to be naïve-like |
| 3a and 3b | Effector-like cell proliferation creating a naïve daughter cell | This may only occur when effector cells can proliferate and when proliferating cells form new less differentiated cells, which we assume to be naïve-like |
| ($\overline{1a}$ or 1c) and $\overline{1d}$ | Differentiation from memory to effector | This may only occur when differentiation between memory-like and effector-like compartments is allowed, and when differentiation is linear from memory-like cells to effector-like cells or it is bidirectional |
| 2b | Differentiation from memory to dysfunction-associated | This may only occur when memory-like cells can upregulate inhibitory receptors |
| (1a or 1c) and $\overline{1d}$ | Differentiation from effector to memory | This may only occur when differentiation between memory-like and effector-like compartments is allowed, and when differentiation is linear from effector-like cells to memory-like cells or it is bidirectional |
| $\overline{3a}$ | Memory-like cell proliferation creating two memory-like daughter cells | This may only occur when proliferating cells do not form new less differentiated (naïve-like) cells |
| 2c | Differentiation from effector-like to dysfunction-associated | This may only occur when effector cells can upregulate inhibitory receptors |
| 1g | Differentiation from dysfunction-associated to effector | This may only occur when dysfunction-associated cells may downregulate inhibitory receptors and back-differentiate to effector-like cells |
| $\overline{3a}$ and 3b | Effector-like cell proliferation creating two effector-like daughter cells | This may only occur when effector-like cells proliferate and proliferating cells do not form new less differentiated (naïve-like) cells |
| 3d | Effector-like cell death | This may only occur when effector-like cells can die |
| 3e | Dysfunction-associated cell death | This may only occur when dysfunction-associated cells can die |

assumed that a feature is off if it is not addressed in the article). Note that we assumed that feature 1e

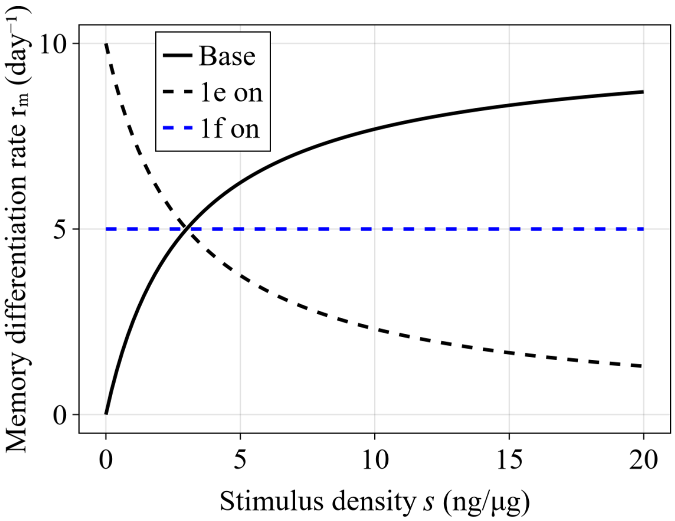


Figure S4.2: **Rate of differentiation to memory-like cells under different feature selections.** Values of the rate of differentiation to memory-like cells $r_{\mathrm{m}}$ as stimulus density $s$ increases under the cases in Eq (5). The base case is when features 1e and 1f are off (first case), 1e on is the second case, and 1f on is the third case. Here, for the purpose of illustration, $r_{\mathrm{m}}^{\max} = 10\,\mathrm{day}^{-1}$ and $s_{\mathrm{m}} = 5\,\mathrm{ng}/\mu\mathrm{g}$.

Table S4.2: **Feature selections in case studies.** Selections (on or off) for each feature to generate the pathways and models in case study 1 [6] and case study 2 [7].

| Case study | Feature selection | | | | | | | | | | | | | | |
|---|---|---|---|---|---|---|---|---|---|---|---|---|---|---|---|
| | 1a | 1b | 1c | 1d | 1e | 1f | 1g | 2a | 2b | 2c | 3a | 3b | 3c | 3d | 3e |
| 1 | on | off | off | off | off | off | on | off | off | on | off | off | off | off | on |
| 2 | on | on | off | off | off | off | off | off | on | off | off | off | off | on | off |

is off in the first case study for the purpose of demonstration (this pathway seems to describe *ex vivo* expansion more accurately without feature 1e on), despite Table 2 of the main text indicating that we interpreted implicit support for feature 1e being on in the pathway suggested by Wherry [6]. The key features of this pathway are present for the demonstration (Fig 3) in the main text.

As an example, we now detail the derivation of the mathematical model that we simulated for the first case study. Feature 1a is on, so the condition "1a or 1b or 1c" is true and the second term in Eq (1) is present in this model. Since we also have that feature 1b is off, both the conditions "$\overline{\text{1a}}$ and $\overline{\text{1c}}$" and "1b" are false so the condition for the first term in Eq (1) is false and the term does not appear in this model. Features 2a, 3a, 3b and 3c are off, so all other conditions indicated in Eq (1) are false and their corresponding terms are also not in this model. We repeat this process for all conditions in Eqs (2)–(4) to derive the full model:

$$\begin{aligned}
\frac{\mathrm{d}N}{\mathrm{d}t} &= -r_{\mathrm{e}}(s)N, \\
\frac{\mathrm{d}M}{\mathrm{d}t} &= r_{\mathrm{m}}(s)E + \mu M, \\
\frac{\mathrm{d}E}{\mathrm{d}t} &= r_{\mathrm{e}}(s)N - r_{\mathrm{m}}(s)E - r_{\mathrm{d}}(s)E + r_{\mathrm{e}}(s)D, \\
\frac{\mathrm{d}D}{\mathrm{d}t} &= r_{\mathrm{d}}(s)E - r_{\mathrm{e}}(s)D - \lambda D,
\end{aligned}$$

where

$$r_{\mathrm{m}}(s) = \frac{r_{\mathrm{m}}^{\max} s}{s_{\mathrm{m}} + s},$$
$$r_{\mathrm{e}}(s) = \frac{r_{\mathrm{e}}^{\max} s}{s_{\mathrm{e}} + s},$$
$$r_{\mathrm{d}}(s) = \frac{r_{\mathrm{d}}^{\max} s}{s_{\mathrm{d}} + s},$$

which are identified by evaluating the conditions in Eqs (5)–(7) with features 1e and 1f off. Specifically, "$\overline{1\mathrm{e}}$" and "not(1f and (1a or 1b or 1c))" are both true since features 1e and 1f off, so the first case is selected for both $r_{\mathrm{m}}$ (Eq (5)) and $r_{\mathrm{e}}$ (Eq (6)).
