## Supplementary Text 5 for "A computational investigation of pathways for *ex vivo* differentiation and dysfunction of healthy and patient T cells"

ORCIDs: MSL, 0009-0009-6633-2044; YB, 0000-0001-9901-7782; PRB, 0000-0003-3962-5393; ALJ, 0000-0001-9103-7092

October 19, 2026

### S5 Text. Details for solving the mathematical models and parameter estimation

Without computationally efficient fitting algorithm, fitting all 7,840 models to data with a ten-start approach requires significant computation time. Improving computational efficiency is especially important if this method is extended to simulate more cell subtypes or to consider more pathway features. The Broyden–Fletcher–Goldfarb–Shanno optimisation algorithm involves using the gradient of the loss function to iteratively step towards a local minimum in parameter space. In Julia, packages may be used to numerically estimate the gradient, but this is computationally inefficient. Optim.jl [1] supports the use of an analytically derived gradient of the loss function to greatly improve computation time (in our case, this is approximately ten times faster than using numerically derived gradients for the whole fitting procedure). Here we detail the derivation of an analytic expression for the gradient of the loss function that we use to fit our models to a given data set (Eq (8) of the main text), which is reiterated here in an equivalent form:

$$\mathcal{L}(\boldsymbol{p}) = \frac{1}{|S|}\sum_{s\in S}\|\boldsymbol{u}(T;\boldsymbol{p},s) - \widetilde{\boldsymbol{u}}_s\|^2, \tag{1}$$

where $S$ is the set of stimulus densities used to expand T cells in the experiments (with $|S|$ elements), $\boldsymbol{u}(T;\boldsymbol{p},s) = [N(T;\boldsymbol{p},s), M(T;\boldsymbol{p},s), E(T;\boldsymbol{p},s), D(T;\boldsymbol{p},s)]^{\mathrm{T}}$ contains the solution of the specified ordinary differential equation model under the parameters $\boldsymbol{p} = [r_{\mathrm{m}}^{\max}, r_{\mathrm{e}}^{\max}, r_{\mathrm{d}}^{\max}, s_{\mathrm{m}}, s_{\mathrm{e}}, s_{\mathrm{d}}, \mu]^{\mathrm{T}} \in \mathbb{R}^7$ and stimulus density $s$ at time $T = 8$ days, and $\widetilde{\boldsymbol{u}}_s$ is the vector containing corresponding Day 8 cell subpopulations in the data for stimulus density $s$. The solution to the model $\boldsymbol{u}(t;\boldsymbol{p},s)$ is found by solving the system of ordinary differential equations

$$\frac{\mathrm{d}\boldsymbol{u}}{\mathrm{d}t} = A(\boldsymbol{p},s)\boldsymbol{u},$$

where $A(\boldsymbol{p},s)$ is the coefficient matrix containing the appropriate rates of differentiation, upregulation of inhibitory receptors, proliferation, and death for the given model (this changes depending on the combination of feature selections). Under the initial condition $\boldsymbol{u_0}$, which is fixed during parameter fitting and matches the average initial number of healthy donor or patient-derived T cells in the experiments, the solution to this system at time $T$ is

$$\boldsymbol{u}(T;\boldsymbol{p},s) = e^{A(\boldsymbol{p},s)T}\boldsymbol{u_0}. \tag{2}$$

The matrix exponential is numerically computed using LinearAlgebra.jl in Julia, which implements the scaling and squaring method, a computationally efficient method of estimating matrix exponentials [2].

The gradient of $\mathcal{L}(\boldsymbol{p})$ in Eq (1) with respect to all parameters is $\boldsymbol{\nabla_p}\mathcal{L} = \left[\frac{\partial \mathcal{L}}{\partial p_1}, \ldots, \frac{\partial \mathcal{L}}{\partial p_7}\right]^{\mathrm{T}}$, where $p_i$ is the $i$th parameter in $\boldsymbol{p}$. The $i$th entry in $\boldsymbol{\nabla_p}\mathcal{L}$ is

$$\begin{aligned}\frac{\partial \mathcal{L}}{\partial p_i} &= \frac{1}{|S|}\sum_{s\in S}\frac{\partial}{\partial p_i}\|\boldsymbol{u}(T;\boldsymbol{p},s) - \widetilde{\boldsymbol{u}}_s\|^2 \\ &= \frac{2}{|S|}\sum_{s\in S}\frac{\partial \boldsymbol{u}(T;\boldsymbol{p},s)}{\partial p_i}\cdot(\boldsymbol{u}(T;\boldsymbol{p},s) - \widetilde{\boldsymbol{u}}_s),\end{aligned} \tag{3}$$

where the second equality is by application of the chain rule.

Since a matrix and its derivative do not necessarily commute [3], the chain rule cannot be used directly to calculate the partial derivatives of the matrix exponential in Eq (2). Instead, by the limit definition of the partial derivative, these derivatives are

$$\frac{\partial \boldsymbol{u}(T;\boldsymbol{p},s)}{\partial p_i} = \lim_{h\to 0}\frac{e^{A(\boldsymbol{p}+h\boldsymbol{e_i},s)T} - e^{A(\boldsymbol{p},s)T}}{h}\boldsymbol{u_0},$$

where $\boldsymbol{e_i}$ is the $i$th standard basis vector in parameter space. Taylor expanding $A(\boldsymbol{p}+h\boldsymbol{e_i},s)$ about $p$ gives

$$\begin{aligned}\frac{\partial \boldsymbol{u}(T;\boldsymbol{p},s)}{\partial p_i} &= \lim_{h\to 0}\frac{e^{A(\boldsymbol{p},s)T + h\frac{\partial A}{\partial p_i}T + \mathcal{O}(h^2)} - e^{A(\boldsymbol{p},s)T}}{h}\boldsymbol{u_0} \\ &= L\left(A(\boldsymbol{p},s)T, \frac{\partial A}{\partial p_i}(\boldsymbol{p},s)T\right)\boldsymbol{u_0},\end{aligned} \tag{4}$$

given that the higher order terms vanish in the limit as $h \to 0$ (which may be seen by considering the Taylor series of the matrix exponential), and where $L(A, E) = \lim_{h\to 0} \frac{e^{A+hE} - e^A}{h}$ is the Fréchet derivative of the matrix exponential $e^A$ in the direction of the matrix $E$ [2]. This Fréchet derivative may be obtained by exponentiating an appropriate block matrix and extracting the upper-right block of the resulting block matrix;

$$\exp\left(\begin{bmatrix} A & E \\ 0 & A \end{bmatrix}\right) = \begin{bmatrix} e^A & L(A, E) \\ 0 & e^A \end{bmatrix}.$$

Therefore, by exponentiating the block matrix $\begin{bmatrix} AT & \frac{\partial A}{\partial p_i} T \\ 0 & AT \end{bmatrix}$ using Julia, the Fréchet derivative in Eq (4) and then the partial derivative in Eq (3) may be estimated for each parameter $p_i$. Optim.jl then uses these partial derivatives of $\mathcal{L}(\boldsymbol{p})$ to minimise the loss function in Eq (1).
